# Genetic background shapes AI-predicted variant effects

**DOI:** 10.64898/2026.04.04.715328

**Authors:** Brian M. Schilder, Zhihan Liu, John J. Desmarais, David Laub, Fahimeh Rahimi, Palash Sethi, Lucas A. Pereira, Mengyi Sun, Justin B. Kinney, David M. McCandlish, Juannan Zhou, Peter K. Koo

**Affiliations:** Simons Center for Quantitative Biology, Cold Spring Harbor Laboratory, Cold Spring Harbor, NY, USA; Division of Biomedical Informatics, University of California San Diego, La Jolla, CA, USA; Department of Biology, University of Florida, Gainesville, FL, USA

## Abstract

Predicting the consequences of genetic variants remains a major goal in biomedicine. Conventional approaches typically assess single-nucleotide variants in the context of a single reference genome, without accounting for genetic diversity that can modulate variant effects. Here we introduce the personalized variant effect predictor (pVEP) framework, which quantifies how genetic background across thousands of human genomes from globally diverse populations shapes computational predictions of clinical variant effects. Across deep learning models spanning protein structure, splicing, and noncoding regulation, pVEP reveals that many clinical variants exhibit heterogeneous predicted effects across haplotypes, with the same variant predicted to be pathogenic in some genetic backgrounds and benign in others. We find support for underlying molecular mechanisms, including shifts in predicted protein contacts and changes in splice-site recognition. Overall, personalized genomic context emerges as a systematically underappreciated variable in variant annotation and clinical interpretation, with particular implications for genetically diverse populations.

---

For decades, the biomedical community has sought to assign genetic variants as “benign” or “pathogenic” through a combination of case studies, population-level investigations, experimental evidence, and computational predictions^1–3^. Despite considerable efforts, over half of all entries in ClinVar remain as variants of unknown significance (VUS) or have conflicting labels^4,5^. Even for those with more confident annotations, large-scale population studies have shown that supposedly pathogenic variants often do not cause disease, reflecting incomplete penetrance^6–9^. Although several factors likely contribute to this discrepancy, it is notable that virtually all variant annotation approaches adopt a *one-variant-one-effect* conceptual framework^10^. Variant effect predictors (VEPs) are a class of computational methods designed to quantify the molecular and phenotypic consequences of genetic polymorphisms^2^. Recent advances in deep learning now enable high-throughput variant effect prediction using models trained on evolutionary sequence diversity^11–17^ or supervised sequence-to-function mappings^18–24^. VEP scores are typically computed as the difference in predicted fitness or molecular function between a reference sequence with and without the variant of interest introduced *in silico*. However, this implicitly assumes that the human reference genome is sufficiently representative of all humans^10,25^, despite the fact that each personal genome differs from the reference by roughly 4 to 5 million genetic variants^26,27^.

Here, we introduce the personalized variant effect predictor (pVEP) framework, which quantifies how predicted variant effects are modulated by genetic background. To demonstrate its utility, we applied pVEP to background sequences derived from 3,891 diverse human genomes from the International Genome Sample Resource^28,29^. Using deep learning models for protein fitness scoring, splice-site prediction, and RNA coverage prediction from DNA sequence, we predicted the effects of *∼*85,000 missense, splice-altering, and UTR variants across millions of haplotypes (Fig. 1). By applying explainable artificial intelligence (XAI) methods, we found support for background-dependent interactions between clinical variants and local haplotypes, revealing interpretable mechanisms including shifts in protein contacts and altered splice-site recognition. Together, these results highlight the limitations of conventional reference-only VEP scores and motivate a broader shift toward personalized, context-aware variant interpretation.

**Figure 1.**
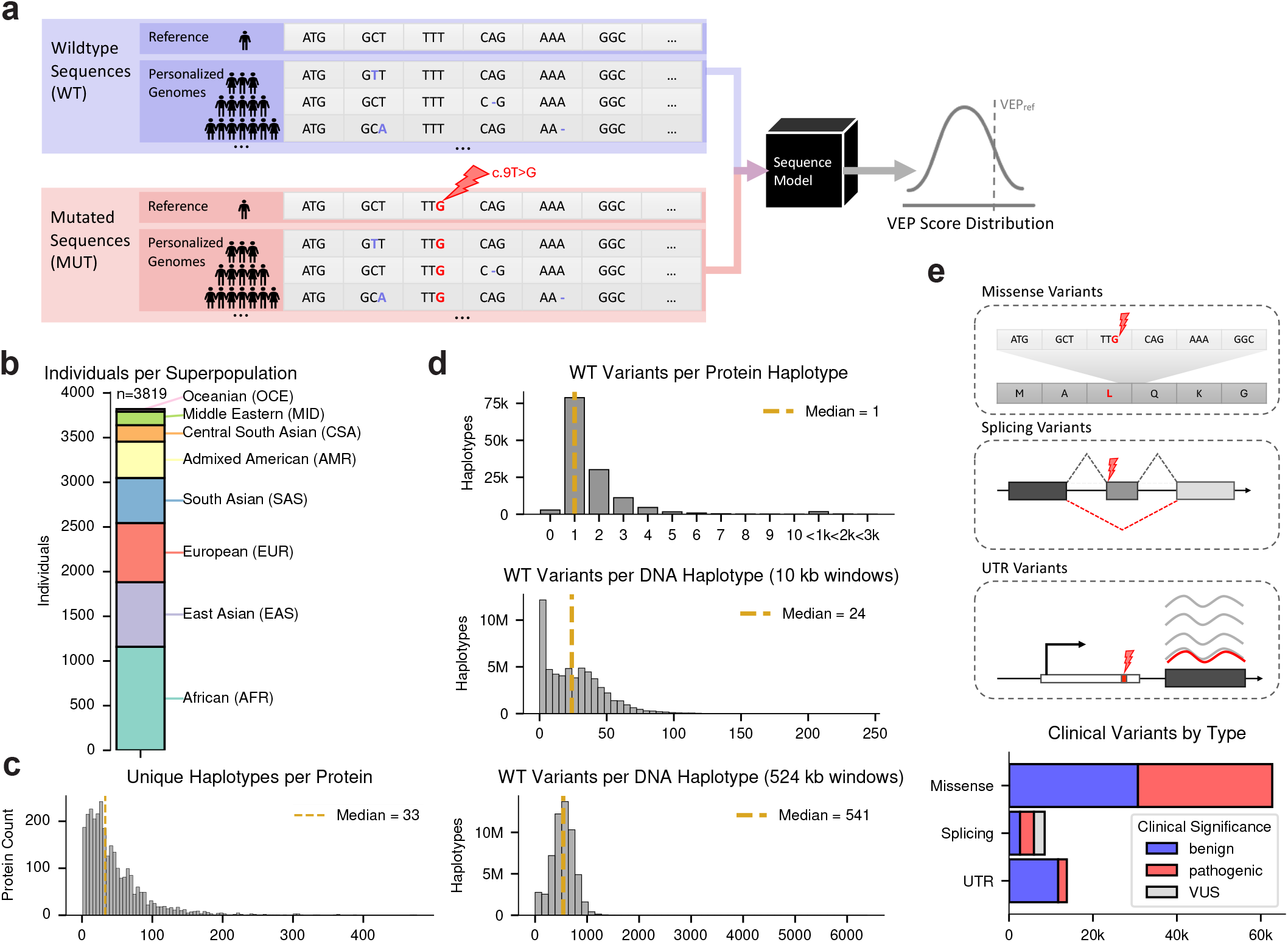
Overview of the pVEP framework and variant datasets. **a**, Schematic of the pVEP framework. For each clinical variant, model predictions are computed across reference and personalized background sequences with (MUT) and without (WT) the clinical variant introduced, yielding a distribution of predicted variant effects with the reference-based VEP score (*VEP*_ref_) marked for comparison. **b**, Bar chart of individuals per ancestral superpopulation across the 1000 Genomes Project and Human Genome Diversity Project (*n* = 3,819 individuals). **c**, Histogram of unique haplotypes per protein across all transcripts analyzed (*n* = 2,847 transcripts). **d**, Histograms of background variants per haplotype for protein sequences in ProteinGym (*top, n* = 132,721 unique haplotypes), 10 kb DNA windows centered on splicing variants in SpliceVarDB (*middle, n* = 64,846,620 unique haplotypes), and 524 kb DNA windows centered on 5^*′*^/3^*′*^ UTR variants from ClinVar (*bottom, n* = 105,182,898 unique haplotypes). Counts *<*10 are binned for visualization purposes. **e**, Schematics of the three clinical variant classes explored and bar plot of variant counts per class (missense: *n* = 62,727, splicing: *n* = 8,490, and UTR: *n* = 13,771), colored by ClinVar pathogenicity label.

## Results

### Population-derived haplotypes define diverse genetic backgrounds for variant effect prediction

To apply pVEP across diverse genetic backgrounds, we assembled personalized haplotypes from 3,819 genomes spanning the 1000 Genomes Project^29^ and the Human Genome Diversity Project^28^ (Fig. 1b), providing a representative sampling of globally diverse human genomes. Here, we use the term *haplotypes* to refer to unique, contiguous DNA or protein sequences from phased human genomes. At the protein level, we reconstructed 132,721 distinct haplotypes across 2,847 transcripts derived from population variants including both single-nucleotide substitutions and indels (Fig. 1c). Most protein haplotypes differed from the reference by a single variant (59%), with the remainder carrying multiple substitutions or indels (Fig. 1d, *top*). As expected from global population structure, protein haplotype diversity was highest in African-ancestry genomes (27% unique; 39% observed at least once), followed by East Asian (17% unique) and European (11% unique) genomes (Supplementary Fig. 1)^28–30^. By contrast, virtually all RNA/DNA sequence haplotypes, defined as 10 kb/524 kb windows centered on each clinical variant, were unique within their loci owing to the larger window sizes and weaker selective constraints in noncoding regions (*N*_haplotypes_ = *N*_variants_*× N*_individuals_ *×* ploidy). This scaling of diversity from compact exons to expansive noncoding intervals defines a broad, population-derived range of genetic backgrounds for our downstream analyses.

To evaluate how clinical variant effects vary across diverse genetic backgrounds, we applied pVEP to three classes of clinical variants: 62,727 ClinVar-annotated missense variants from the ProteinGym benchmark^31^, 8,490 splice-altering variants from SpliceVarDB^32^, and 13,771 ClinVar variants in untranslated regions (UTRs) (Fig. 1e). Each variant was introduced into all compatible personalized haplotypes, and VEP scores were computed as the difference in model predictions between the personalized sequence with and without the clinical variant. This procedure yielded a distribution of VEP scores for each variant across various backgrounds (see Methods). For each molecular level, VEP scores were computed using: multiple ESM models^11,13,14^ for proteins, SpliceAI^19^ for RNA splicing, and Flashzoi^18^, an efficient derivative of Borzoi^21^ for investigating regulatory variants in DNA sequences (Supplementary Table 1).

### Background-dependent missense VEP scores reveal complex pathogenicity landscapes

At the protein level, pathogenic variants were consistently assigned lower predicted fitness than benign ones (Fig. 2a, Supplementary Fig. 1c). To assess how well reference-based predictions represent the full pVEP score distribution, we computed the percentile of the reference-based VEP score within each variant’s population VEP score distribution (Fig. 2b). For a given variant, a 50th percentile would indicate that the reference-based VEP score ranked towards the middle of the full pVEP score distribution and is thus more representative, whereas values closer to 0% or 100% indicate the reference sequence is poorly representative. We then binned the “representativeness scores” into deciles along the *x-axis* (Q1 to Q10, according to per-variant mean pVEP) and quintiles along the *y-axis*. This revealed that the reference-based VEP score fell at extreme percentiles for a substantial fraction of clinical variants, systematically underestimating the pathogenicity of clinically severe missense variants. This deviation grew more pronounced with increasing sequence diversity (Supplementary Fig. 2b–d), indicating that reference-genome predictions are often poor representatives of the population-level VEP score distribution.

**Figure 2.**
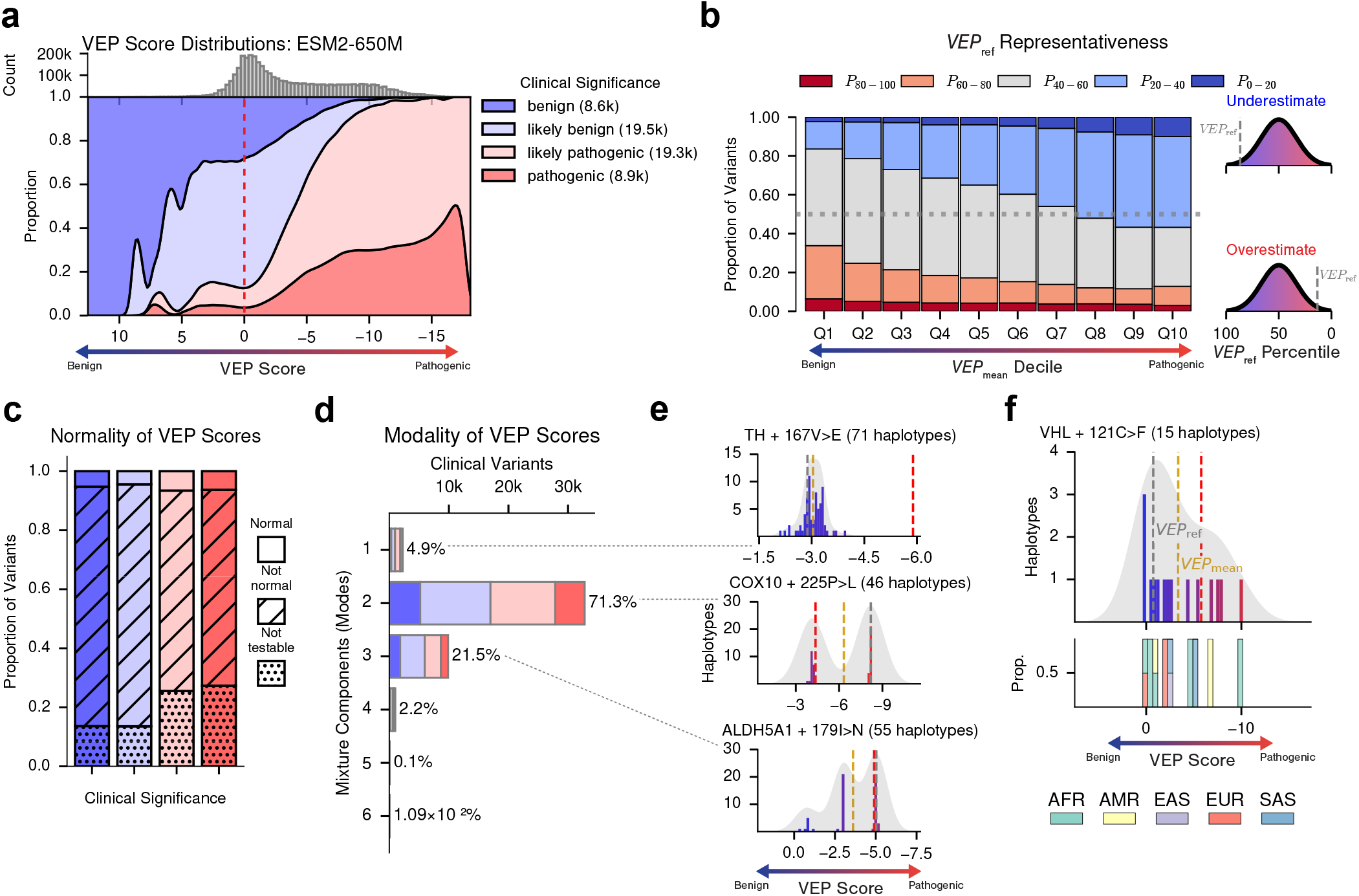
pVEP reveals heterogeneous predicted responses to clinical missense variants. **a**, Filled density plot of ESM2-650M reference-based and personalized VEP scores across all missense variants, stratified by ClinVar pathogenicity (*n* = 62,727: *∼* 8, 600 benign, *∼* 19, 500 likely benign, *∼* 19, 300 likely pathogenic, and *∼* 8, 900 pathogenic variants). Negative values (*rightwards*) indicate more pathogenic effects. Vertical dashed line (*red*) indicates benign/pathogenic threshold. **b**, Stacked bar plot showing the percentile bin (*fill color*) of reference-based VEP score (*VEP*_ref_) relative to the full distribution average (*VEP*_mean_) at each *VEP*_mean_ decile. A greater proportion of red bars indicates *VEP*_ref_ tends to overestimate pathogenicity, whereas a greater proportion of blue bars indicates underestimation. The gray horizontal dashed line demarcates 50% of the variants in that VEP Decile. **c**, Stacked barplot showing the proportion of clinical variants whose ESM2-650M VEP scores follow a normal distribution (D’Agostino-Pearson omnibus test, FDR *>* 0.05), non-normal distribution (FDR *<* 0.05,) or could not be tested (*n <* 20 protein haplotypes), grouped by ClinVar pathogenicity. **d**, Bar plot of the estimated number of modes per clinical variant VEP score distribution, inferred from a variational Bayes Gaussian mixture model. **e**, Haplotype-level VEP score distributions for three missense variants with high inter-haplotype variability: TH 167V*>*E (*n* = 71 haplotypes, *top*), COX10 225P*>*L (*n* = 46, *middle*), and ALDH5A1 179I*>*N (*n* = 55, *bottom*). Vertical dashed lines indicate *VEP*_mean_ (*gold*), *VEP*_ref_ (*grey*), and the empirically inferred benign/pathogenic boundary (*red*). **f**, Haplotype-level ESM2-650M VEP score distribution for VHL 121C*>*F (*n* = 15 haplotypes). Vertical dashed lines indicate *VEP*_mean_ (*gold*) and *VEP*_ref_ (*grey*). Bar plots below show the proportion of individuals from each ancestry per VEP score bin. AFR=African, AMR=Admixed American, EAS=East Asian, EUR=European, SAS=South Asian.

Across thousands of clinical missense variants observed in at least 20 unique haplotypes, only *∼*5% of population VEP score distributions followed a unimodal normal distribution (Fig. 2c). The majority instead showed multimodal structure: fitting a variational Bayes Gaussian mixture model^33,34^ revealed two or more distinct peaks in more than 95% of variants (Fig. 2d,e). Among all loci analyzed, VHL showed striking background-dependent variability. VHL variants cause Von Hippel-Lindau disease, a multisystem disorder predisposing to diverse cancers^35,36^. The VHL 121C*>*F variant, which is annotated in ClinVar as being pathogenic, displayed highly variable effects across haplotypes, ranging from highly pathogenic to completely benign (Fig. 2f), with several effects being ancestry-specific. Notably, an African-ancestry haplotype produced the most pathogenic response (VEP score *∼−* 10), while the reference-based VEP score alone would classify this variant as benign, illustrating how reference-based predictions provide an incomplete view of variant effects across genetic backgrounds.

### Protein haplotypes modulate the pathogenicity of structurally proximal cancer risk variants

Given the extensive molecular and clinical characterization of *BRCA1*^37–41^, we used it as a model system to dissect the determinants of background-dependent pVEP scores. We developed a post hoc interpretability approach, termed a *variant sensitization map*, that uses an interpretable linear model to quantify how each background variant shifts the predicted effect of each clinical variant across haplotypes (see Methods; Fig. 3a). Intuitively, a background variant that consistently makes a clinical variant more damaging will have a large positive coefficient, while one that buffers its effect will have a large negative coefficient.

**Figure 3.**
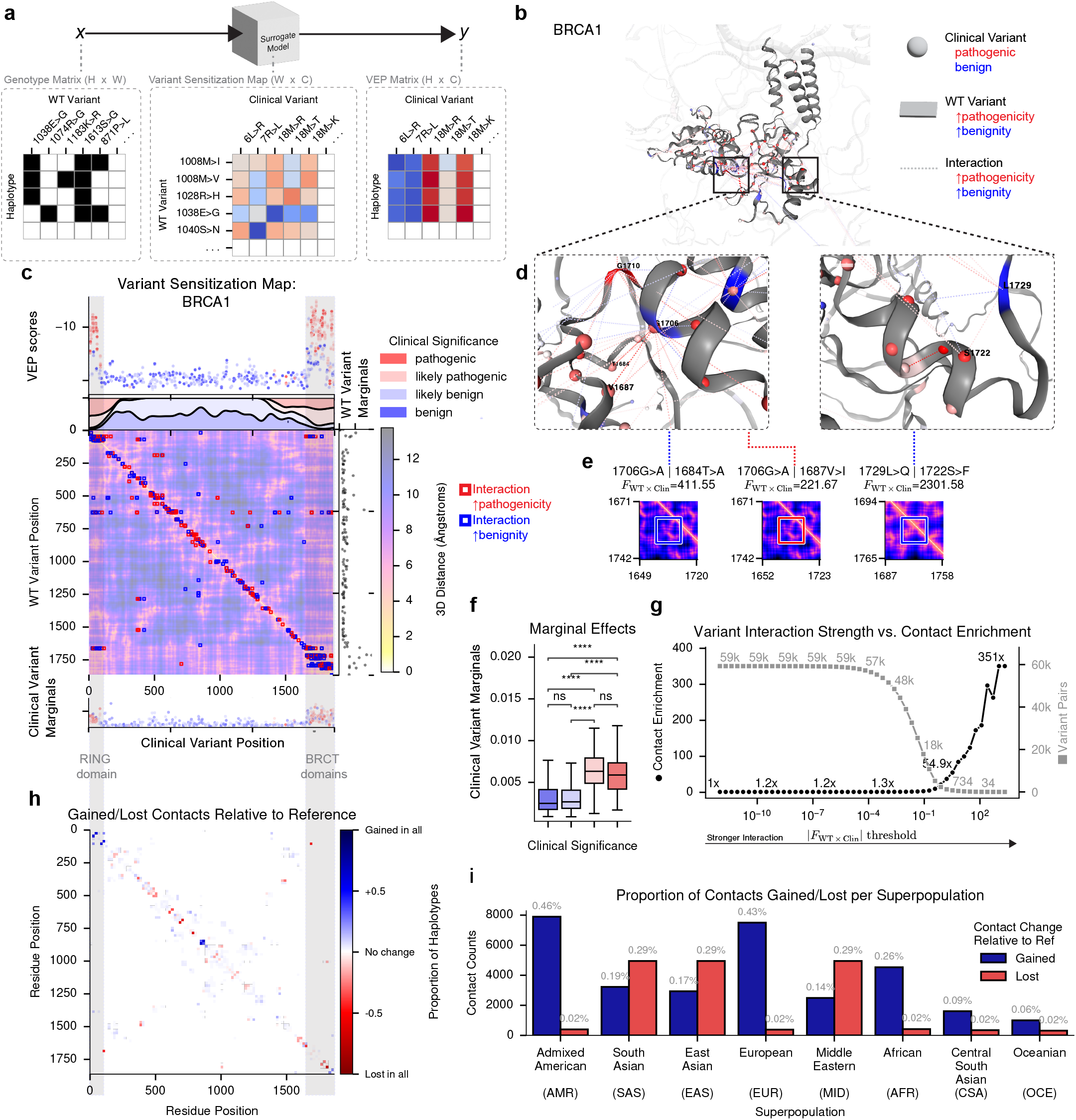
Background variants modify the effects of BRCA1 clinical variants in close structural proximity. **a**, Surrogate modeling takes as input a binary matrix of background variant presence across haplotypes (*left*) and predicts continuous clinical variant VEP scores (*right*), yielding a variant sensitization map of inferred joint effects (*center*). **b**, Predicted 3D structure of BRCA1 from AlphaFold2 (AF2). **c**, Variant sensitization map (inferred from ESM2 VEP scores) overlaid on the AF2-predicted BRCA1 distance map, showing all 652 significant non-additive interactions (*center heatmap*; *p <* 0.05). Red boxes indicate variant pairs where a background variant (*y-axis*) increases predicted pathogenicity of a clinical variant (*x-axis*). Pathogenic clinical variants (*top scatter/density*) are concentrated in the RING and BRCT domains (*grey vertical bars*), recapitulated by surrogate model marginal effects for clinical (*bottom*) and background variants (*right*). **d**, 3D structural visualization of interaction pairs: background variant 1706G*>*A with clinical variants 1684T*>*A and 1687V*>*I (*left*), and 1729L*>*Q with 1722S*>*F (*FDR* = 1.43 *×*10^*−*13^; *right*). Spheres indicate clinical variants colored by *VEP*_mean_ (*red*: pathogenic, *blue*: benign); ribbon color and dashed lines indicate the background variant’s effect on pathogenicity (*red*: *↑, blue*: *↓, grey*: no data). Labels indicate reference amino acid and residue position. **e**, Zoomed-in AF2 distance maps for the four variant pairs with the strongest interaction coefficients in BRCA1. **f**, Boxplot of clinical variant marginals grouped by ClinVar pathogenicity; pathogenic variants have higher marginals than benign ones (two-sided Mann–Whitney *U* tests: *U* = 1355–2302, *p* = 1.33×10^*−*21^ to *×*8.33 10^*−*13^, *n*_benign*/*likely benign_ = 137–167, *n*_pathogenic*/*likely pathogenic_ = 57–95; ns: *p ≥* .05; ****: *p≤* 0.0001). **g**, Observed-to-expected enrichment of AF2 residue contacts among variant pairs at each interaction threshold, with variant pair counts on the secondary axis. **h**, Residue-by-residue heatmap showing the proportion of haplotypes (*n* = 126) gaining (*blue*) or losing (*red*) AF2-predicted contacts (*<* 8 Å) relative to the reference. **i**, Bar plot of reference contacts gained (*blue*) or lost (*red*) in population-weighted contact maps stratified by superpopulation; labels show percentage gained/lost relative to the reference.

Fitting this model across 132 background and 456 clinical variants in 144 haplotypes explained a large proportion of the variance (*R*^2^ = 0.83; mean-squared error = 0.00016). Approximately 1% of background *−*clinical variant pairs were significantly non-additive (interaction test: FDR *<* 0.05; see Supplementary Methods), and the strongest interactions were concentrated at three-dimensional residue contacts predicted by AlphaFold2^16^, particularly within the RING and BRCT domains where most pathogenic BRCA1 variants cluster (Fig. 3b,c). The most pronounced interaction involved the rare BRCT substitution 1729L*>*Q, which is benign in isolation but synergistically amplifies pathogenicity of the breast cancer risk variant 1722S*>*F^37,42–44^ (Fig. 3d, e), a pairwise interaction not, to our knowledge, previously reported. Another BRCA1 WT variant, 1706G*>*A, can either increase or decrease pathogenicity depending on the other variants present (1687V*>*I and 1684T*>*A, respectively). More broadly, pathogenic clinical variants showed markedly stronger marginal effects (calculated as mean absolute joint effect per variant) than benign variants (Fig. 3f). Stronger interactions were progressively more enriched for structural contacts, reaching 351-fold enrichment at the highest thresholds (Fig. 3g).

Applying AlphaFold2 across all 126 BRCA1 haplotypes revealed subtle but consistent background-dependent structural differences: small fractions of residue pairs gained (0.41%) or lost (0.03%) contacts relative to the reference in at least one haplotype (Fig. 3h). This variability was not explained by AlphaFold2 prediction confidence, as measured by the predicted Local Distance Difference Test (pLDDT; Supplementary Fig. 3). Superpopulation-specific contact maps revealed that Admixed American and European haplotypes showed the most contact gains relative to the reference, while South Asian, East Asian, and Middle Eastern haplotypes showed the greatest losses; African haplotypes displayed comparatively modest structural variability despite the highest overall sequence diversity (Fig. 3i).

Parallel analyses in TP53 (26 haplotypes) recapitulated the key trends: interaction strength enriched for protein contacts, and pathogenic variants showed stronger marginal effects than benign ones (Supplementary Fig. 4). Approximately 2.3% of background *−*clinical variant pairs showed evidence of significant non-additivity, including the background variant 215S*>*R which buffered multiple pathogenic alleles within the DNA-binding domain of TP53. Unlike BRCA1, TP53 showed minimal superpopulation-level structural variability, except in East Asian haplotypes, which exhibited a markedly elevated fraction of lost contacts, though whether such losses are functionally consequential remains unclear^45,46^. Together, these findings suggest that background-dependent VEP scores may in part reflect structural reconfiguration of the protein, pointing to a plausible mechanistic basis for context-dependent pathogenicity across human populations.

### Background variation modulates splicing variant effects

We next examined the extent that genetic background modulates predicted effects at the RNA level, focusing on splice-altering mutations. Across SpliceAI-based VEP scores for curated splice-altering variants, ClinVar-labeled pathogenic variants (*n ≈*2, 600) were predicted to be more spliceogenic (larger positive values) than VUS (*n ≈*2, 500) or benign (*n≈* 3, 300 k) variants (Fig. 4a–b). Pathogenic variants predominantly induced acceptor or donor loss, whereas benign variants more often produced acceptor or donor gains (Fig. 4c), and these patterns remained consistent across *∼*55 million personalized predictions. Predicted splicing effects also varied in their stability across haplotypes. Quantifying variability in SpliceAI-predicted effect classes for each clinical variant revealed greater inter-haplotype variability for benign variants and VUS than for pathogenic variants (Fig. 4d), consistent with the expectation that strongly deleterious splice disruptions yield more uniform outcomes^47^.

**Figure 4.**
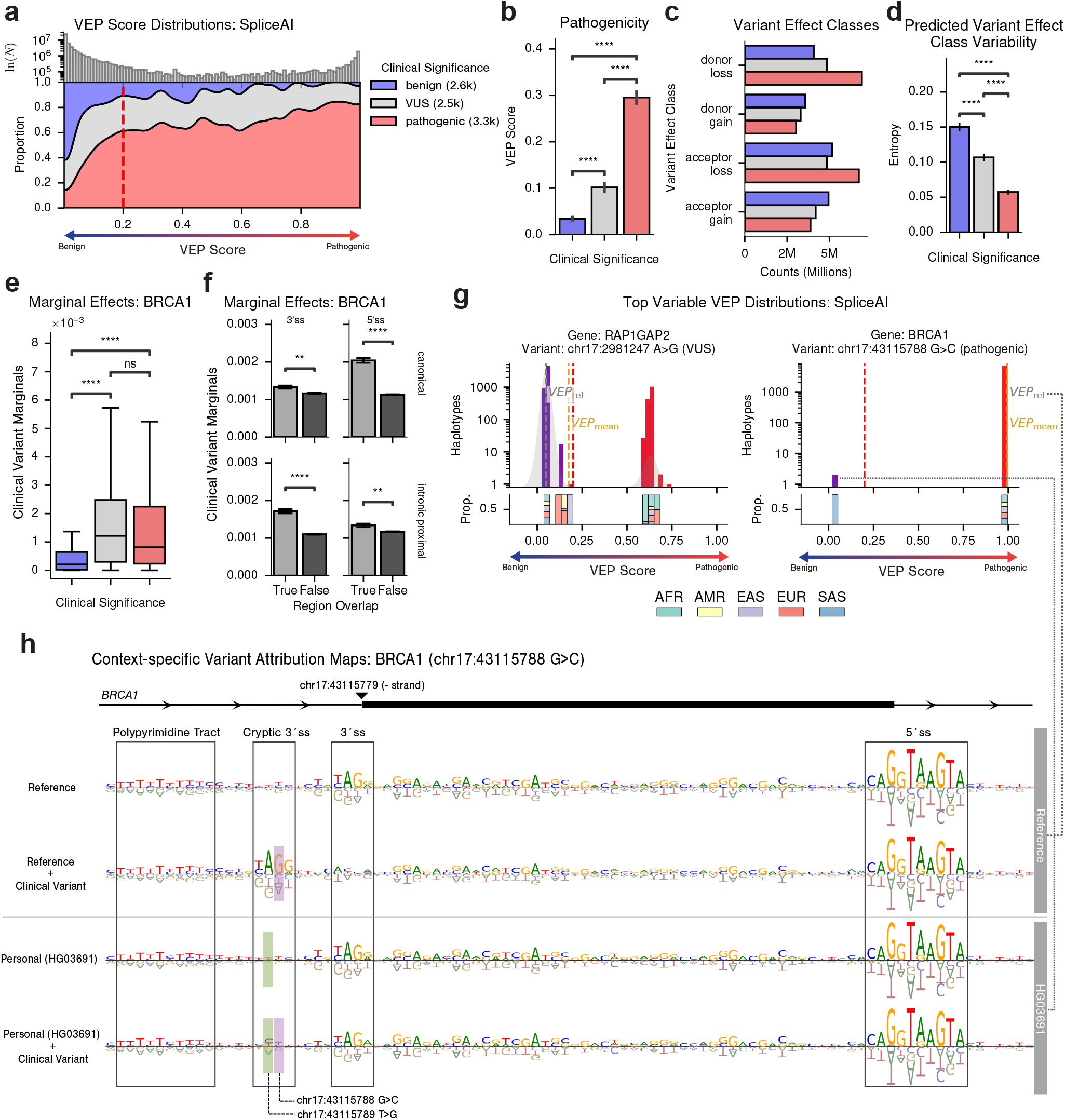
pVEP reveals background-dependent predicted effects for clinical splicing variants. **a**, Filled density plot of all SpliceAI reference-based and personalized VEP scores per clinical variant (x-axis: 0 = no predicted splice disruption, 1 = complete disruption), stratified by ClinVar pathogenicity (*n* = 8.5 k: 2.6 k benign, 2.5 k VUS, 3.3 k pathogenic). The red vertical line indicates spliceogenicity threshold. **b**, Bar plot of pVEP scores grouped by ClinVar pathogenicity label, showing pathogenic variants and VUS have high predicted spliceogenicity than benign ones (two-sided Mann-Whitney U test: *U* = 2.16 10^6^ – 2.72 *×*10^6^, *p* = 2.03 *×*10^*−*114^ to 2.51*×* 10^*−*243^). ns: *p ≥* .05; **: *p <* 0.01; ****: *p <* 0.0001). **c**, Bar plot of counts of SpliceAI-predicted variant effect classes (donor/acceptor gain or loss), colored by clinical significance. **d**, Bar plot of Shannon entropy of SpliceAI-predicted effect classes across haplotypes, grouped by pathogenicity labels. Pathogenic variants and VUS have significantly higher entropy than benign variants (two-sided independent-samples *t*-tests: *t* = 5.52–14.35, *p* = 6.2 *×*10^*−*46^ to 3.6 *×*10^*−*8^) **e**, Boxplot of surrogate model marginal effects per clinical variant, colored by ClinVar significance, showing pathogenic variants and VUS have higher marginal effects than benign variants (two-sided Mann-Whitney *U* tests: *p <* 0.0001). **f**, Bar plot of clinical variant marginals grouped by whether the clinical variant overlaps a splicing-relevant region: 3^*′*^ splice-site (ss), 5^*′*^ss, canonical, or intronic proximal. Variants within canonical and intronic proximal 3^*′*^/5^*′*^ss displayed greater effects than others (two-sided Mann-Whitney *U* tests: *p <* 0.001). **g**, VEP score distributions across haplotypes for chr17:2981247 A*>*G (*RAP1GAP2*, VUS) and chr17:43115788 G*>*C (*BRCA1*, pathogenic). Upper panels show log-scale haplotype frequency with spliceogenicity (*red*) and reference (*grey*) thresholds marked. Lower panels show superpopulation composition per VEP score bin. **h**, Attribution maps for four genomic context around *BRCA1* chr17:43115788 G*>*C: reference and HG03691 haplotype, each with and without the clinical variant. Attribution maps given by Integrated Gradients highlight sequence-level determinants of 3^*′*^ splice-site choice. The clinical variant is highlighted in *purple* and the compensatory background variant in *green*. This variant induced a strong splicing change in all haplotypes (max Δ = 0.996) except for the HG03691-specific haplotype (max Δ = 0.0195).

To characterize the sequence determinants of these background-dependent splicing effects, we extended our variant sensitization method to relate background variant presence to SpliceAI-based VEP scores in *BRCA1*. As in the protein analysis, both pathogenic clinical variants and VUS exhibited significantly higher marginal effects than benign variants (Fig. 4e). Furthermore, clinical variants within core splicing machinery (canonical and intronic proximal 3^*′*^/5^*′*^ss) showed significantly greater marginal effects than others (Fig. 4f). We also explicitly tested splicing variants for non-additive interactions. Background−clinical variant pairs within the same exon exhibited stronger interactions than those across distant exons (Supplementary Fig. 5d), consistent with known splice-site mechanisms, and background variants within the same proximal exonic 5^*′*^ss as the clinical variant produced the strongest modulation of VEP scores (Supplementary Fig. 5e), reflecting the narrow sequence constraints of proximal splice-site elements.

Among all splicing variants, chr17:2981247A*>*G (a VUS in the gene *RAP1GAP2*) and chr17:43115788 G*>*C (a *BRCA1* variant labeled as pathogenic in ClinVar; Fig. 4g) displayed the strongest haplotype-dependent effects. In most haplotypes, the latter *BRCA1* mutation induced a strong predicted splicing effect, which attribution analysis identified as a cryptic acceptor gain (Fig. 4h). However, in two South Asian-specific haplotypes, the predicted effect fell below the splice-altering threshold. Further attribution analysis revealed that these haplotypes carry a counteractive A*>*C substitution that disrupts the cryptic AG dinucleotide, preventing aberrant splice-site activation (Fig. 4h). This suggests a plausible molecular mechanism by which a background variant can attenuate the impact of an otherwise pathogenic clinical mutation.

As orthogonal experimental support, we compared SpliceAI predictions against a massively parallel splicing assay (MPSA) comprising *∼* 30,000 *BRCA2* minigene variants with double-mutant perturbations^48,49^. Predicted splice-site scores correlated with experimentally measured splicing ratios and recovered both additive and pairwise epistatic coefficients (Supplementary Fig. 5). These results support the capacity of SpliceAI to capture pairwise epistatic interactions between variants, lending credibility to the background-dependent variant effects identified using pVEP across diverse haplotypes.

### Background variation modulates UTR variant effects

We next applied pVEP to noncoding regulatory regions, computing reference- and population-based VEP scores for *∼* 14,000 ClinVar variants in 5^*′*^ and 3^*′*^ UTRs using Flashzoi within 524 kb windows. Pathogenic variants showed significantly higher predicted regulatory impact than benign ones in both 3^*′*^ UTR, and 5^*′*^ UTR regions (Fig. 5a–b). Within the benign subset, 5^*′*^ UTR variants displayed modestly stronger and more variable predicted effects than 3^*′*^ variants (Fig. 5b–c). Ranking variants by predicted inter-haplotype variability revealed substantial heterogeneity (Fig. 5d). The pathogenic 5^*′*^ UTR variant in *CTNS* (chr17:3655058 C*>*T, underlying nephropathic cystinosis) and *MLH1* (chr3:37004474 G*>*C, associated with Lynch Syndrome) showed more than an order-of-magnitude difference in predicted activity across haplotypes. In both cases, the reference-based VEP scores markedly underestimated the population average. Conversely, two benign-labeled 3^*′*^ UTR variants in *FGD4* (chr12:32645731 G*>*A) and ARSA (chr22:50623794 G*>*C) exhibited clearly multimodal distributions in which many haplotypes crossed into the pathogenic range. Track-level Flashzoi predictions likewise revealed substantial inter-haplotype variability, with some variants showing minimal effects in the reference sequence but strong predicted perturbations across many haplotypes (Supplementary Fig. 6–7). These results show that background-dependent VEP score heterogeneity extends across all molecular layers examined, from protein-coding to noncoding regulatory regions.

**Figure 5.**
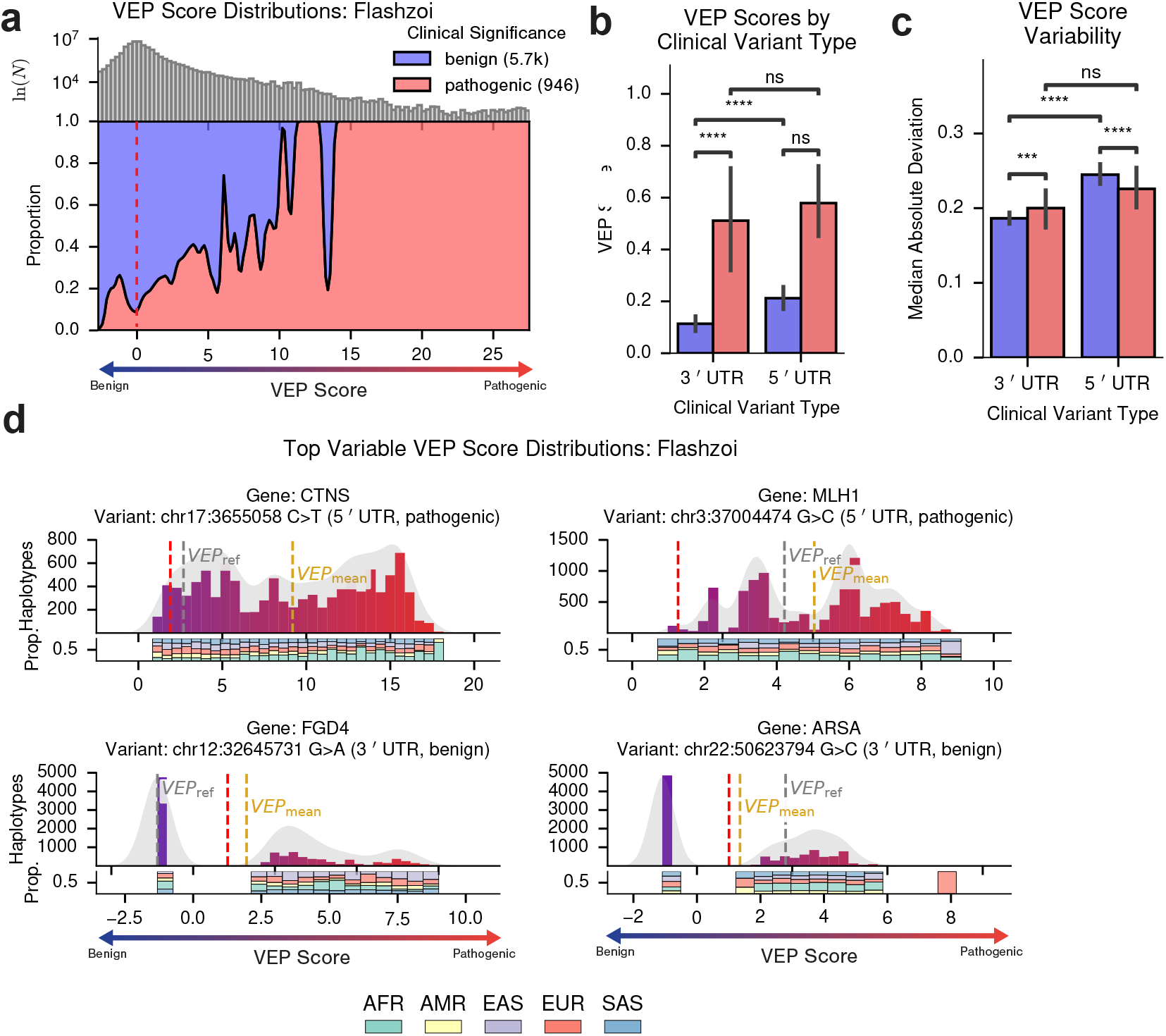
pVEP reveals background-dependent predicted effects for clinical UTR variants. **a**, Filled density plot of Flashzoi average pVEP scores (*VEP*_mean_) per clinical variant, stratified by ClinVar pathogenicity. Vertical dashed lines indicate the reference-based VEP score (*VEP*_ref_; *grey*) and the empirically inferred benign/pathogenic boundary (*red*). **b**, Bar plot of *VEP*_mean_ grouped by UTR type (5^*′*^/3^*′*^) and ClinVar pathogenicity label. Pathogenic-labeled variants had significantly greater predicted pathogenicity than benign variants (two-sided Mann–Whitney *U* tests: 3^*′*^ UTR benign vs.pathogenic, *U* = 4.42*×*10^5^, *p* = 1.3*×*10^*−*7^; 5^*′*^ UTR, *U* = 4.65 10^5^, *p* = 0.016). ns: *p ≥*.05; ****: *p≤* 0.0001. **c**, Bar plot of variability (median absolute deviation) of Flashzoi VEP score distributions across haplotypes, grouped by UTR type and ClinVar pathogenicity label. Benign variants within 5^*′*^ UTR regions showed greater variability in predicted effects than those witihn 3^*′*^ regions (two-sided Mann-Whitney *U* test: *p <* 0.0001). **d**, Haplotype-level VEP score distributions for the four clinical UTR variants with the greatest variability across haplotypes. Vertical dashed lines indicate *VEP*_mean_ (*gold*), *VEP*_ref_ (*grey*), and the empirically inferred benign/pathogenic boundary (*red*), derived from gene-specific variational Bayes Gaussian mixture models. Below each histogram, bar plots show the proportion of haplotypes from each ancestry per VEP bin.

### Population-averaged VEP scores improve correspondence with functional annotations

Given that reference-based VEP scores reflect only a single genetic background, whereas pVEP reveals substantial heterogeneity across backgrounds, we reasoned that averaging pVEP scores across haplotypes would provide a more representative estimate of variant effect. To assess this, we computed Spearman correlations between the reference-based VEP score or averaged pVEP scores against 95 established variant annotations from Ensembl VEP^3^ (spanning evolutionary conservation, population frequency, and orthogonal pathogenicity). Across all three models, the averaged pVEP scores yielded stronger statistically significant correlations for the majority of annotations, with the most pronounced improvements observed for Flashzoi (Supplementary Fig. 8a–c). The largest gains occurred for annotations aligned with each model’s primary biological domain: mutfunc and AlphaMissense^50,51^ for protein-level predictions, MaxEntScan^52^ for splicing, and dbscSNV spliceogenicity scores for UTR effects^53^.

Across all tested models, Flashzoi showed the strongest and most consistent improvements in annotation correlations when comparing averaged pVEP scores to reference-based VEP scores (95.9% win rate across all annotations). In particular, averaged pVEP scores from Flashzoi strengthened the expected negative relationship between pathogenicity and allele frequency relative to reference-based VEP scores alone (Supplementary Fig. 8d). Computing superpopulation-specific pVEP score averages and matching them to allele frequencies from the corresponding superpopulation further reinforced this relationship (Supplementary Fig. 9). These results indicate that pVEP-derived population-averaged predictions show broader and stronger correspondence with established variant annotations than single reference-based estimates.

## Discussion

Variant interpretation is central to both clinical genetics and biological research^2^. Although population-scale sequencing has revealed extensive human genetic diversity^6,27–29^, variant interpretation workflows remain largely reference-centric^2,10^. Modern deep learning-based VEP approaches have inherited this convention. Despite being trained on evolutionarily or population-diverse sequence corpora, variants are typically evaluated only in the context of the reference genome^11–24^, thereby implicitly assuming that predicted effects generalize across individuals. To address this gap, we introduced pVEP, a framework that systematically quantifies how genetic background modulates predicted variant effects across thousands of globally diverse haplotypes.

By applying pVEP across deep learning models for protein, splicing, and regulatory sequences, we found that the predicted effects of clinical variants are often sensitive to genetic background, leading to broad and often multimodal distributions. For some variants, predictions remained stable across backgrounds, indicating that reference estimates are sometimes representative. However, in many other cases, the reference genome poorly represented the broader distribution of predicted effects across sampled human haplotypes. In this sense, background dependence is itself an important property of variant effect estimates.

We found indirect support for molecular mechanisms underlying some of the observed background-dependent effects. In proteins, the strongest modifiers clustered near predicted residue contacts. In splicing, modifiers localized to canonical splice-relevant regions and included compensatory variants that redirect cryptic splice-site choice. Although these observations remain indirect, they suggest that at least some of the observed background dependence may reflect biologically meaningful local sequence context.

A recurring pattern was that some haplotypes, often more common in particular ancestry groups, produced distinctive predicted effects. For example, some African haplotypes generated the most pathogenic predicted response to the VHL 121C>F variant, whereas some South Asian haplotypes carried a counteractive substitution that suppressed a pathogenic BRCA1 splicing effect. These observations point to a broader equity concern: conventional VEP approaches systematically ignore genetic contexts that differ from the reference genome, which is primarily North/West European and West African^54^, potentially contributing to biases in the clinical interpretation of variants. The *one-variant-one-effect* assumption is therefore not only scientifically limiting, but may also disproportionately affect the populations least represented in existing genomic databases.

These findings should also be interpreted in the context of current limitations in sequence modeling. The accuracy and biological fidelity of sequence models for variant effect prediction remain open questions, particularly in personalized-genome settings. Prior work has shown that models such as Enformer^55^ can struggle to predict molecular phenotypes such as gene expression from personalized genomes^56–58^. We do not resolve those limitations here. Instead, the strongest support in this study comes from prediction settings with more established precedent for variant interpretation, particularly protein function and splicing, where models such as ESM and SpliceAI have shown strong performance and where some of the observed background dependence admits more direct local molecular interpretation. We also note that several newer sequence models emerged after this study was initiated, including Evo 2^59^ and AlphaGenome^20^, and were therefore not evaluated here. Our results should therefore be interpreted not as a comprehensive comparison of the current model landscape, but as evidence that background sensitivity can be substantial even in widely used variant effect prediction settings.

More broadly, this study should be viewed as a proof of concept for the pVEP framework in a globally diverse but still limited cohort. Applying the same framework to substantially larger resources, such as All of Us^60^ or UK Biobank^61^, could enable a more comprehensive assessment of how human genetic background shapes variant effect predictions across a wider spectrum of human diversity.

The extent to which this heterogeneity reflects biological context dependence, model misspecification, or a combination of both remains unresolved in the absence of population-scale experimental ground truth. At present, there are no systematic experimental resources that quantify the molecular or phenotypic consequences of the same variant across diverse human genetic backgrounds, limiting validation, calibration, and clinical translation of VEP tools. Addressing these questions will likely depend on progress across several fronts, including large-scale perturbational assays in genetically diverse cellular or organoid systems to directly measure background-dependent variant effects, cohort-scale studies linking phased genomes to molecular or clinical phenotypes, and benchmarking frameworks that treat variant effects as conditional distributions over genetic backgrounds.

Variants of unknown significance and conflicting annotations currently constitute the majority of ClinVar entries^1,4,5^, and most putatively pathogenic variants have near-zero penetrance in real-world populations^6–9^. Our pVEP framework offers one possible path forward, as ambiguous variant consequences may in some cases reflect sequence context dependence. Establishing these empirical foundations will be essential for determining whether personalized variant effect prediction can support equitable and accurate interpretation at clinical-grade reliability.

## Methods

### Personalized variant effect prediction (pVEP) framework

We developed the pVEP framework to compute variant effect predictions (VEP scores) contextualized to individual genetic backgrounds. Population haplotypes carry diverse background variants but typically lack the focal variant of interest (e.g., ClinVar variants). pVEP *in silico* injects each focal variant into every haplotype, scores the resulting wild-type (WT) and mutant (MUT) sequences with pretrained models, and returns per-haplotype VEP scores from delta between WT and MUT,. This yields a *distribution* of predicted effects across a cohort. The framework comprises complementary DNA and protein pipelines unified by the following workflow:

1. **For each** focal variant *v ∈ V* :
2. **For each** haplotype *h ∈ H* :
3. Extract WT sequence: **s**^wt^ *←* ExtractSeq(*h*, locus(*v*))
4. Inject focal variant: **s**^mut^ *←* InjectVariant(**s**^wt^, alt(*v*))
5. **For each** model *m ∈ M* :
6. VEP_*v,h,m*_ *←* Δ(*m*(**s**^mut^), *m*(**s**^wt^))
7. Collect *{*VEP_*v,h,m*_*}*_*h,m*_ as the personalized VEP score distribution for *v*.

### Protein pipeline

The protein pipeline retrieves population haplotype sequences per transcript via the Ensembl Hap-losaurus API^30^ (`haplosaurus.get_haplotypes`), then injects each focal mutation into every haplotype background (`haplosaurus.add_variant_to_haplotypes`). The entry point (`vep_pipeline`) scores each WT/MUT pair via `ESM_predict`, with per-residue log-likelihood differences computed by `vep_metrics.compute_mt_wt_score`. Results are stored as Parquet files (one per model–protein–haplotype–strategy combination):

~~~
*cohort*}/vep/
   {*model*}/
       {*protein*}/
           {*haplotype*}/
                {*variant_source*}/
                      **wt-marginals.parquet
                            masked-marginals.parquet
                                  pseudo-ppl.parquet**
~~~

### DNA pipeline

The DNA pipeline uses GenVarLoader (GVL)^62^ to load phased haplotypes from population-scale variant call sets and a reference genome. For each focal variant and sample, GVL extracts a genomic window centered on the variant within the sample’s phased haplotype context (`haps_to_seqs`), then constructs MUT sequences by inserting the focal alternate allele. Both haploid copies are scored independently, yielding two VEP estimates per diploid individual. The entry point (`vep_pipeline`) dispatches to model-specific modules, each implementing `load_model()`, `load_tokenizer()`, and `run_vep()`. Results are stored as xarray datasets in compressed zarr format (one store per chromosome):

~~~
*cohort*}/{*variant_set*}/
  **chr1.zarr**
    dims: *site* × *sample* × *ploid* × *metric*
   vars: one array per model
   **chr2.zarr**
   …
~~~

### Extensibility

New protein models can be added for any architecture that accepts a protein sequence and returns per-position logits; new DNA models can be integrated by implementing the three-function interface (`load_model`/`load_tokenizer`/`run_vep`). Input focal variants (e.g., ClinVar variants) are specified as PLINK/VCF/BCF files or site tables; population data as Haplosaurus-compatible cohorts (protein) or phased PLINK/VCF/BCF genotype files (DNA), allowing direct use of any biobank with phased sequencing data. Downstream analyses—surrogate modeling, attribution, and epistasis testing—operate on the standardized output coordinates and are model-agnostic. All code is publicly available (see Code Availability section).

### Population cohorts and haplotype construction

Personalized haplotypes were derived from 3,819 phased genomes spanning two publicly available cohorts: the 1000 Genomes Project (1KGP) Phase 3^29^ and the Human Genome Diversity Project (HGDP)^28^, together capturing broad global ancestry diversity across seven superpopulations (African, Admixed American, Central/South Asian, East Asian, European, Middle Eastern, and Oceanian). Superpopulation assignments for 1KGP samples followed the standard IGSR metadata labels. For HGDP samples, which are annotated with fine-grained population labels (e.g., Yoruba, French, Han), we mapped each population to its corresponding superpopulation using the HGDP–IGSR population metadata^28^.

#### Protein haplotypes

Personalized protein sequences were reconstructed using the Haplosaurus module of Ensembl VEP^30^. For 1KGP samples, haplotypes were retrieved via the Ensembl REST API. For HGDP samples, Haplosaurus was run locally on phased, merged autosomal VCFs from the International Genome Sample Resource (IGSR), aligned to the GRCh38 reference genome. Only variants passing quality filters (INFO score ≥ 0.9) were retained; both single-nucleotide variants (SNVs) and short insertion-deletion variants (indels) were included. This yielded 132,721 distinct protein haplotypes across 2,847 transcripts.

#### DNA haplotypes

Personalized DNA sequences were constructed from the same 1KGP and HGDP VCFs using `genvarloader`^63^. Each clinical variant was centered within a genomic window, with window sizes matched to the context requirements of each sequence model: 10,101 nt for SpliceAI predictions (a 101 nt scoring window with 5,000 bp flanking on each side) and 524,288 bp for Flashzoi predictions. Full data accession details are provided in the **Data Availability** section.

### Variant datasets

Three classes of clinical variants were analyzed, matched to the appropriate molecular model.

#### Missense variants

Clinical missense variants were obtained from ProteinGym v1^31^, which compiles ClinVar-annotated^1^ pathogenic and benign substitutions into a standardized benchmark. Because ProteinGym’s original workflow is reference-centric, we developed a custom Python pipeline to inject each variant into all compatible personalized protein haplotypes, correcting residue indices to account for coding indels and preserve positional accuracy across haplotypes. This yielded 62,727 missense variants for analysis.

#### Splicing variants

Splice-altering variants were curated from ClinVar and SpliceVarDB^32^ (both accessed March 2025), retaining only SNVs annotated as benign or pathogenic. Intersecting the two sources yielded 8,490 validated variants. Each variant was annotated to a canonical genomic subregion (canonical splice site, intronic proximal, exonic, or core splice site) using MANE Select transcripts^64^.

#### UTR variants

5^*′*^ and 3^*′*^ UTR variants were curated from the GRCh38-formatted ClinVar VCF (accessed May 2025), filtering for SNVs with review status ≥2 stars and molecular consequences annotated as `5_prime_UTR_variant` or `3_prime_UTR_variant` using `genoray.VCF`^63^. This yielded 13,771 variants labeled as benign or pathogenic.

### Protein variant effect prediction

VEP scores for missense variants were computed using models from the Evolutionary Scale Modeling (ESM) suite, including ESM1b, ESM1v, ESM2 (35M, 650M, and 15B parameters), ESM3 (1.4B), and ESMC (600M)^11,13–15^ (Supplementary Table 1). For all models, we applied the masked marginals scoring strategy recommended by the ESM authors^13^, which infers per-residue log-likelihood differences between mutant and wild-type sequences using the masked language modeling objective. Log-likelihood differences (Δlog *L*) were computed via `torch.nn.functional.log_softmax` and interpreted as predicted relative sequence fitness. We used ESM2-650M for primary analyses, as it achieves near-state-of-the-art zero-shot fitness prediction performance while remaining computationally tractable for population-scale inference across millions of personalized haplotypes.

### VEP distribution analysis

#### Modality

For each clinical variant with ≥ 20 unique protein haplotypes, the resulting distribution of VEP scores was assessed for normality and modality. Normality was tested using the D’Agostino-Pearson omnibus test (`scipy.stats.normaltest`), with significance defined at a false discovery rate (FDR) *>* 0.05. The number of modes in each distribution was estimated using a variational Bayes Gaussian mixture model^33,34^ (`sklearn.mixture.BayesianGaussianMixture`, maximum 50 components).

#### Reference representativeness

To quantify how representative reference-based predictions (*VEP*_ref_) were of population-derived distributions, we computed the percentile rank of *VEP*_ref_ within the full set of pVEP scores for each variant (Fig. 2b, 2). We used percentile-based comparisons rather than z-scores due to the non-normality of these distributions. Variants were stratified by mean pathogenicity quantiles (*VEP*_mean_) and visualized as stacked histograms to show over-or underestimation trends of *VEP*_ref_ relative to the population distribution.

### Surrogate modeling and variant sensitization maps

To identify which background variants modulate the predicted pathogenicity of clinical variants, we fitted regularized linear surrogate models relating background haplotype composition to VEP scores.

#### Protein surrogate models

For each protein, a binary matrix **X** (haplotypes × background variants) encoded the presence or absence of each wild-type (WT) variant in each haplotype, and a continuous matrix **Y** (haplotypes× clinical variants) contained ESM2-650M VEP scores for each haplotype–clinical variant pair (Fig. 3a). Ridge regression (`sklearn.linear_model.Ridge`, *α* = 1.0, random seed = 42) was used to fit:

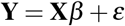

The resulting coefficient matrix *β* constitutes the *variant sensitization map*: |*β*_*i j*_|quantifies the magnitude of the joint effect between background variant *i* and clinical variant *j*, and the sign of *β*_*i j*_ encodes its direction (negative values indicating increased predicted pathogenicity). Model fit was evaluated using *R*^2^, mean squared error (MSE), and related metrics from `sklearn.metrics`. The same surrogate modeling pipeline was applied to both BRCA1 (126 haplotypes, 1,788 clinical variants) and TP53 (26 haplotypes, 189 clinical variants) using ESM2-650M.

#### Splicing surrogate models

For splicing analyses, where the relevant genomic context is localized, a separate single-target surrogate model was fitted per clinical variant rather than the multi-target approach used for proteins. The feature matrix **X** was restricted to background WT variants within a ±5 kb window of the focal variant, ensuring that each model captures only variants falling within the receptive field of SpliceAI during VEP score inference. The response variable *y* was the SpliceAI VEP score (maximum absolute delta score) across haplotypes for that single clinical variant. Ridge regression was applied with the same hyperparameters (*α* = 1.0, random seed = 42).

Full mathematical details of both surrogate modeling approaches are provided in the Supplementary Methods.

### Non-additive interaction testing

To test whether the joint effect of a WT–clinical variant pair deviates from the sum of their individual effects, we compared two nested Ridge regression models for each pair: an additive model (*y* = *β*_0_ + *β*_1_*g*) and an interaction model (*y* = *β*_0_ + *β*_1_*g* + *β*_2_*g*·*d*), where *g* indicates WT variant presence and *d* is the deviation of the observed joint effect from the expected additive sum. Significance was assessed via F-test (*p <* 0.05; `scipy.stats.f`), restricted to pairs with ≥ 10 haplotypes and representation of both WT-present and WT-absent groups. For protein analyses, individual effects were estimated from the multi-target surrogate model (averaged across all clinical sites per protein); for splicing analyses, individual effects were estimated per clinical variant from the single-target surrogate models, restricted to WT variants within the same ±5 kb window. Full details are provided in the Supplementary Methods.

Full details of both interaction testing procedures are provided in the Supplementary Methods.

### Structural contact enrichment analysis

To assess whether non-additive interactions between WT and clinical variant pairs (*β*_WT_×_Clin_) were enriched at three-dimensional residue contacts, we used AlphaFold2 (AF2)^16^ contact maps predicted from the reference sequence. Contacts were defined as residue pairs with predicted C*α*–C*α* distance *<*8 Å. Enrichment at a given interaction strength threshold *t* was defined as the ratio of observed to expected overlap between significant joint effects (|*β*_*i j*_|*> t*) and predicted contacts:

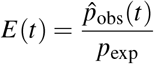

where 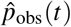is the proportion of variant pairs exceeding threshold *t* that are also in contact, and *p*_exp_ is the overall contact frequency for that protein. Significance at each interaction threshold was assessed with a binomial test (`scipy.stats.binomtest`) and *p <* 0.05 was considered significant.

### Personalized protein structure analysis

AF2^16^ was run independently on each unique protein haplotype for BRCA1 (126 haplotypes) and TP53 (26 haplotypes). Residue-residue contact maps were binarized at 8 Å. For each haplotype, contacts were compared to the reference contact map to identify gained and lost contacts. To test whether predicted structural variability across haplotypes could be explained by model uncertainty, we computed the Pearson correlation between per-position pairwise contact variance and predicted Local Distance Difference Test (pLDDT) scores.

To examine ancestry-associated structural variation, superpopulation-specific contact maps were derived as frequency-weighted averages over all haplotypes, with each haplotype weighted by its observed frequency within the target superpopulation. This avoids hard ancestry assignments and ensures that haplotypes shared across populations contribute proportionally to each group’s structural profile. Haplotypes common in a given ancestry therefore contribute more to that superpopulation’s map, while haplotypes rare or absent in that ancestry contribute little or nothing.

### Splicing variant effect prediction

SpliceAI-10K^19^ was applied to both reference and variant-inserted personalized haplotypes using a 10,101 nt input window. For each sequence, donor and acceptor gain and loss probabilities were computed across the central 101 nt scoring window, yielding four delta scores per variant. The maximum absolute delta score was taken as the VEP score for each haplotype, reflecting the strongest predicted perturbation to splicing.

To quantify inter-haplotype consistency in predicted outcomes, each haplotype was assigned its predominant effect class (donor gain, donor loss, acceptor gain, or acceptor loss), and Shannon entropy^65^ was computed over the resulting probability distribution of classes across haplotypes for each clinical variant. A score of zero indicates that all haplotypes share the same predicted consequence, while higher values reflect increasing background-dependent variability in splicing outcome.

### Splicing attribution analysis

To characterize the sequence-level basis for context-dependent splicing outcomes, integrated gradients^66^ were computed for the BRCA1 variant chr17:43115788 G*>*C and a South Asian haplotype carrying a compensatory A*>*C substitution. Attribution maps were calculated using SEAM^67^ with 25 integration steps, using the summed SpliceAI acceptor probability across the 101 nt scoring window as the target function. Attribution scores were visualized as sequence logos using `LogoMaker`^68^.

### Splicing epistasis benchmarking

To validate SpliceAI’s ability to recover local pairwise interactions, we analyzed a massively parallel splicing assay (MPSA) dataset comprising ∼30, 000 variants at the 9-nt core 5^*′*^ splice site of a *BRCA2* minigene^48^. Following the approach of Tareen et al.^49^, we fitted a MAVE-NN linear model with additive and pairwise terms to SpliceAI-predicted donor logits and compared the inferred coefficients to those derived from experimental exon inclusion measurements. Agreement was assessed using Pearson correlations for both additive and epistatic coefficients.

### UTR variant effect prediction

UTR VEP scores were computed using Flashzoi^18^, a computationally efficient extension of Borzoi^21^, which predicts genome-wide RNA coverage tracks from 524 kb input sequences. Following the Borzoi convention, per-variant effects were quantified as a Coverage Ratio (COVR) summarizing the maximum predicted fold-change in RNA coverage across all output tracks:

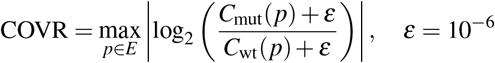

where *C*_mut_(*p*) and *C*_wt_(*p*) are the predicted coverage values at genomic position *p* for the mutant and wild-type sequences, respectively, and the maximum is taken over all positions in the genomic window *E*. Track-wise Shannon entropy was computed using `scipy.stats.entropy` to identify regulatory signals with the greatest inter-haplotype variability.

### Benchmarking population-averaged VEP scores

To assess whether population-derived predictions better capture known variant properties, Spearman correlations were computed between VEP scores (reference-derived *VEP*_ref_ or population-averaged *VEP*_mean_) and 95 genome-wide variant annotations from Ensembl VEP^3^, spanning evolutionary conservation, population allele frequencies, and pathogenicity scores from orthogonal computational tools. Only annotations for which both *VEP*_ref_ and *VEP*_mean_ were significantly correlated (*p <* 0.05) were included in the comparison. An improvement “win rate” was defined as the proportion of annotation features for which |*r*_mean_|*>*| *r*_ref_ | (strict inequality; ties were counted as neither a win nor a loss).

The relationship between Flashzoi VEP scores and population allele frequencies was modeled using a three-parameter logistic function, *f* (*x*) = *L/*(1+exp(*™k*(*x™x*_0_))), with parameters estimated by nonlinear least squares (`scipy.optimize.curve_fit`).

To further examine ancestry-specific concordance, superpopulation-averaged VEP scores were matched to allele frequencies from the corresponding superpopulation and compared against population-agnostic and reference-only estimates. Model fit was assessed using *R*^2^ and an F-test for significance, with 95% confidence intervals computed from the covariance matrix of estimated parameters.

### Computational resources

All protein VEP scores were computed on 4×NVIDIA A100 80 GB Graphics Processing Units (GPUs). SpliceAI predictions were run on a single NVIDIA H100 80 GB GPU. Flashzoi predictions (524 kb input windows) were run on 20 ×NVIDIA H100 80 GB GPUs in parallel to mitigate the longer inference time and greater number of variants. AlphaFold2 structure predictions were run on 2 ×NVIDIA A100 GPUs using ColabFold^69^. All analyses used Python 3.12.9 with PyTorch 2.x and TensorFlow 2.x backends.

### Statistical analyses and software

All analyses were performed in Python 3.12.9. Statistical tests included the D’Agostino-Pearson omnibus test, two-sided Mann– Whitney U test, two-sided *t*-test, binomial test, and F-test, implemented via `scipy.stats`. Multiple testing corrections used the Benjamini–Hochberg FDR procedure. Linear models were fitted using `scikit-learn`. All source code is publicly available (see **Code Availability**).

## Data Availability

1KGP VCFs are publicly available at: https://ftp.1000genomes.ebi.ac.uk/vol1/ftp/data_collections/1000_genomes_project/release/20190312_biallelic_SNV_and_INDEL/

HGDP VCFs are publicly available at: https://ngs.sanger.ac.uk/production/hgdp/hgdp_wgs.20190516/statphase/

The GRCh38 reference genome FASTA file is publicly available at: https://ftp.1000genomes.ebi.ac.uk/vol1/ftp/technical/reference/GRCh38_reference_genome/GRCh38_full_analysis_set_plus_decoy_hla.fa

ProteinGym variants can be downloaded from: https://proteingym.org/

SpliceVarDB variants can be downloaded from: https://compbio.ccia.org.au/splicevardb/

UTR variants were curated directly from the GRCh38-formatted ClinVar VCF (downloaded May 19, 2025): https://ftp.ncbi.nlm.nih.gov/pub/clinvar/vcf_GRCh38/clinvar.vcf.gz

All pVEP and analysis results generated in this study are publicly accessible on Zenodo: https://doi.org/10.5281/zenodo.18452090

## Code Availability

Code for the Ensembl VEP tool (which includes Haplosaurus) is available on GitHub:

https://github.com/Ensembl/ensembl-vep

Code for the protein sequence models is available on GitHub and Zenodo:

https://github.com/bschilder/VEP_protein

https://doi.org/10.5281/zenodo.18890914

Code for the DNA sequence models is available on GitHub and Zenodo:

https://github.com/bschilder/VEP_DNA

https://doi.org/10.5281/zenodo.18890852

Source code for `GenVarLoader` is available on GitHub:

https://github.com/mcvickerlab/GenVarLoader

Source code for ColabFold, and its extension for running on local compute (localcolabfold), are available on GitHub

https://github.com/sokrypton/colabfold

https://github.com/YoshitakaMo/localcolabfold

## Acknowledgments

This work was supported in part by: the Simons Center for Quantitative Biology at Cold Spring Harbor Laboratory as well as NIH grants: R01HG012131, R01GM149921, R35GM133613, and R35GM154908. CSHL Cancer Center Developmental Funds partly supported this work from the Cancer Center Support Grant P30CA045508, and computations were performed using equipment supported by NIH grant S10OD028632.

We would also like to thank Mafalda Dias, Jonathan Frazer, Charles Pugh, Federico Billeci, Carlos Martí, and Evan Seitz for their invaluable feedback and guidance throughout this project.

## Author Contributions

BMS, PKK and JZ conceptualized and designed the study. BMS coded and executed the analyses for the ESM and Flashzoi VEP pipelines, the personalized AF2 prediction pipeline, surrogate modeling for all modalities, and all other analyses except where otherwise specified. ZL and JD coded and executed the SpliceAI VEP pipeline. ZL performed the SpliceAI epistasis and variant attribution analyses. DL developed and supported analyses with GenVarLoader. FR, PS and LP ran exploratory analyses and provided assistance as needed. MS ran Haplosaurus on the Human Genome Diversity Project. BMS wrote the initial draft, and all authors helped edit. PKK, BMS, and JZ primarily supervised the project, with additional supervisory support by DMM and JBK.

## Competing Interests

The authors have no competing interests to declare.

## Notes

https://doi.org/10.5281/zenodo.18452090

https://ftp.1000genomes.ebi.ac.uk/vol1/ftp/data_collections/1000_genomes_project/release/20190312_biallelic_SNV_and_INDEL/

https://ngs.sanger.ac.uk/production/hgdp/hgdp_wgs.20190516/statphase/

https://ftp.1000genomes.ebi.ac.uk/vol1/ftp/technical/reference/GRCh38_reference_genome/GRCh38_full_analysis_set_plus_decoy_hla.fa

https://proteingym.org/

https://compbio.ccia.org.au/splicevardb/

https://ftp.ncbi.nlm.nih.gov/pub/clinvar/vcf_GRCh38/clinvar.vcf.gz

### Competing Interest Statement

The authors have declared no competing interest.

## References

1. Landrum, M. J. et al. ClinVar: Public archive of interpretations of clinically relevant variants. Nucleic Acids Res. 44, D862–D868, 10.1093/nar/gkv1222 (2016).

2. Riccio, C., Jansen, M. L., Guo, L. & Ziegler, A. Variant effect predictors: a systematic review and practical guide. Hum. Genet. 143, 625–634, 10.1007/s00439-024-02670-5 (2024).

3. McLaren, W. et al. The ensembl variant effect predictor. Genome Biol. 17, 122, 10.1186/s13059-016-0974-4 (2016).

4. Lazareva, T. E., Barbitoff, Y. A., Nasykhova, Y. A. & Glotov, A. S. Major causes of conflicting interpretations of variant pathogenicity in rare disease: A systematic analysis. J. Pers. Medicine 14, 864, 10.3390/jpm14080864 (2024).

5. Bonetti, E., Tini, G. & Mazzarella, L. Accuracy of renovo predictions on variants reclassified over time. J. Transl. Medicine 22, 713, 10.1186/s12967-024-05508-w (2024).

6. Gudmundsson, S. et al. Exploring penetrance of clinically relevant variants in over 800,000 humans from the genome aggregation database. bioRxiv 2024.06.12.593113, 10.1101/2024.06.12.593113 (2024).

7. Wright, C. F. et al. Assessing the pathogenicity, penetrance, and expressivity of putative disease-causing variants in a population setting. The Am. J. Hum. Genet. 104, 275–286, 10.1016/j.ajhg.2018.12.015 (2019).

8. Forrest, I. S. et al. Population-based penetrance of deleterious clinical variants. JAMA 327, 350–359, 10.1001/jama.2021.23686 (2022).

9. Kingdom, R. & Wright, C. F. Incomplete penetrance and variable expressivity: From clinical studies to population cohorts. Front. Genet. 13, 10.3389/fgene.2022.920390 (2022).

10. Ciesielski, T. H., Sirugo, G., Iyengar, S. K. & Williams, S. M. Characterizing the pathogenicity of genetic variants: the consequences of context. npj Genomic Medicine 9, 3, 10.1038/s41525-023-00386-5 (2024).

11. Rives, A. et al. Biological structure and function emerge from scaling unsupervised learning to 250 million protein sequences. Proc. Natl. Acad. Sci. 118, e2016239118, 10.1073/pnas.2016239118 (2021).

12. Riesselman, A. J., Ingraham, J. B. & Marks, D. S. Deep generative models of genetic variation capture the effects of mutations. Nat. Methods 15, 816–822, 10.1038/s41592-018-0138-4 (2018).

13. Meier, J. et al. Language models enable zero-shot prediction of the effects of mutations on protein function, 10.1101/2021.07.09.450648 (2021).

14. Lin, Z. et al. Evolutionary-scale prediction of atomic-level protein structure with a language model. Science 379, 1123–1130, 10.1126/science.ade2574 (2023).

15. Hayes, T. et al. Simulating 500 million years of evolution with a language model. Science 0, eads0018, 10.1126/science.ads0018 (2025).

16. Jumper, J. et al. Highly accurate protein structure prediction with AlphaFold. Nature 596, 583–589, 10.1038/s41586-021-03819-2 (2021).

17. Abramson, J. et al. Accurate structure prediction of biomolecular interactions with AlphaFold 3. Nature 630, 493–500, 10.1038/s41586-024-07487-w (2024).

18. Hingerl, J. C., Karollus, A. & Gagneur, J. Flashzoi: an enhanced borzoi for accelerated genomic analysis. Bioinformatics 41, btaf467, 10.1093/bioinformatics/btaf467 (2025).

19. Jaganathan, K. et al. Predicting splicing from primary sequence with deep learning. Cell 176, 535–548.e24, 10.1016/j.cell.2018.12.015 (2019).

20. Avsec, et al. AlphaGenome: advancing regulatory variant effect prediction with a unified DNA sequence model, 10.1101/2025.06.25.661532 (2025).

21. Linder, J., Srivastava, D., Yuan, H., Agarwal, V. & Kelley, D. R. Predicting RNA-seq coverage from DNA sequence as a unifying model of gene regulation. Nat. Genet. 1–13, 10.1038/s41588-024-02053-6 (2025).

22. Zhou, J. et al. Deep learning sequence-based ab initio prediction of variant effects on expression and disease risk. Nat. Genet. 50, 1171–1179, 10.1038/s41588-018-0160-6 (2018).

23. Zhou, J. & Troyanskaya, O. G. Predicting effects of noncoding variants with deep learning–based sequence model. Nat. Methods 12, 931–934, 10.1038/nmeth.3547 (2015).

24. Kelley, D. R. et al. Sequential regulatory activity prediction across chromosomes with convolutional neural networks. Genome Res. 28, 739–750, 10.1101/gr.227819.117 (2018).

25. Martin, A. R. et al. Human demographic history impacts genetic risk prediction across diverse populations. The Am. J. Hum. Genet. 100, 635–649, 10.1016/j.ajhg.2017.03.004 (2017).

26. Phan, L. et al. The evolution of dbSNP: 25 years of impact in genomic research. Nucleic Acids Res. 53, D925–D931, 10.1093/nar/gkae977 (2025).

27. Trost, B., Loureiro, L. O. & Scherer, S. W. Discovery of genomic variation across a generation. Hum. Mol. Genet. 30, R174–R186, 10.1093/hmg/ddab209 (2021).

28. Bergström, A. et al. Insights into human genetic variation and population history from 929 diverse genomes. Science 367, eaay5012, 10.1126/science.aay5012 (2020).

29. Auton, A. et al. A global reference for human genetic variation. Nature 526, 68–74, 10.1038/nature15393 (2015).

30. Spooner, W. et al. Haplosaurus computes protein haplotypes for use in precision drug design. Nat. Commun. 9, 4128, 10.1038/s41467-018-06542-1 (2018).

31. Notin, P. et al. ProteinGym: Large-scale benchmarks for protein design and fitness prediction. bioRxiv 2023.12.07.570727, 10.1101/2023.12.07.570727 (2023).

32. Sullivan, P. J., Quinn, J. M. W., Wu, W., Pinese, M. & Cowley, M. J. SpliceVarDB: A comprehensive database of experimentally validated human splicing variants. Am. J. Hum. Genet. 111, 2164–2175, 10.1016/j.ajhg.2024.08.002 (2024).

33. Attias, H. A variational bayesian framework for graphical models. Adv. Neural Inf. Process. Syst 12 (2000).

34. Blei, D. M. & Jordan, M. I. Variational inference for dirichlet process mixtures. Bayesian Analysis 1, 121–143, 10.1214/06-BA104 (2006).

35. Gossage, L., Eisen, T. & Maher, E. R. VHL, the story of a tumour suppressor gene. Nat. Rev. Cancer 15, 55–64, 10.1038/nrc3844 (2015).

36. Kaelin, W. G. The von hippel-lindau tumor suppressor protein: roles in cancer and oxygen sensing. Cold Spring Harb. Symp. on Quant. Biol. 70, 159–166, 10.1101/sqb.2005.70.001 (2005).

37. Kinget, L. et al. Multitumor case series of germline BRCA1, BRCA2 and CHEK2-mutated patients responding favorably on immune checkpoint inhibitors. Curr. Oncol. 28, 3227–3239, 10.3390/curroncol28050280 (2021).

38. Lovelock, P. K. et al. Genetic, functional, and histopathological evaluation of two c-terminal BRCA1 missense variants. J. Med. Genet. 43, 74–83, 10.1136/jmg.2005.033258 (2006).

39. Lee, M. S. et al. Comprehensive analysis of missense variations in the BRCT domain of BRCA1 by structural and functional assays. Cancer research 70, 4880–4890, 10.1158/0008-5472.CAN-09-4563 (2010).

40. Zeng, C. et al. Association of pathogenic variants in hereditary cancer genes with multiple diseases. JAMA Oncol. 8, 835–844, 10.1001/jamaoncol.2022.0373 (2022).

41. Golubeva, V. A., Nepomuceno, T. C. & Monteiro, A. N. A. Germline missense variants in BRCA1: New trends and challenges for clinical annotation. Cancers 11, 522, 10.3390/cancers11040522 (2019).

42. Lu, W., Wang, X., Lin, H., Lindor, N. M. & Couch, F. J. Mutation screening of RAD51c in high-risk breast and ovarian cancer families. Fam. Cancer 11, 381–385, 10.1007/s10689-012-9523-9 (2012).

43. Chan, G. H. J. et al. Clinical genetic testing outcome with multi-gene panel in asian patients with multiple primary cancers. Oncotarget 9, 30649–30660, 10.18632/oncotarget.25769 (2018).

44. Reza, M. N. et al. Pathogenic genetic variants from highly connected cancer susceptibility genes confer the loss of structural stability. Sci. Reports 11, 19264, 10.1038/s41598-021-98547-y (2021).

45. Fischer, N. W., Ma, Y.-H. V. & Gariépy, J. Emerging insights into ethnic-specific TP53 germline variants. JNCI J. Natl. Cancer Inst. 115, 1145–1156, 10.1093/jnci/djad106 (2023).

46. Doffe, F. et al. Identification and functional characterization of new missense SNPs in the coding region of the TP53 gene. Cell Death Differ. 28, 1477–1492, 10.1038/s41418-020-00672-0 (2021).

47. Blakes, A. J. M. et al. A systematic analysis of splicing variants identifies new diagnoses in the 100,000 genomes project. Genome Medicine 14, 79, 10.1186/s13073-022-01087-x (2022).

48. Wong, M. S., Kinney, J. B. & Krainer, A. R. Quantitative activity profile and context dependence of all human 5′ splice sites. Mol. Cell 71, 1012–1026.e3, 10.1016/j.molcel.2018.07.033 (2018).

49. Tareen, A. et al. MAVE-NN: learning genotype-phenotype maps from multiplex assays of variant effect. Genome Biol. 23, 98, 10.1186/s13059-022-02661-7 (2022).

50. Slodkowicz, G. & Babu, M. M. From prioritisation to understanding: mechanistic predictions of variant effects. Mol. Syst. Biol. 14, e8741, 10.15252/msb.20188741 (2018).

51. Cheng, J. et al. Accurate proteome-wide missense variant effect prediction with AlphaMissense. Science 381, eadg7492, 10.1126/science.adg7492 (2023).

52. Yeo, G. & Burge, C. B. Maximum entropy modeling of short sequence motifs with applications to RNA splicing signals. J. Comput. Biol. A J. Comput. Mol. Cell Biol. 11, 377–394, 10.1089/1066527041410418 (2004).

53. Jian, X., Boerwinkle, E. & Liu, X. In silico prediction of splice-altering single nucleotide variants in the human genome. Nucleic Acids Res. 42, 13534–13544, 10.1093/nar/gku1206 (2014).

54. Schneider, V. A. et al. Evaluation of GRCh38 and de novo haploid genome assemblies demonstrates the enduring quality of the reference assembly. Genome Res. 27, 849–864, 10.1101/gr.213611.116 (2017).

55. Avsec, et al. Effective gene expression prediction from sequence by integrating long-range interactions. Nat. Methods 18, 1196–1203, 10.1038/s41592-021-01252-x (2021).

56. Huang, C. et al. Personal transcriptome variation is poorly explained by current genomic deep learning models. Nat. Genet. 55, 2056–2059, 10.1038/s41588-023-01574-w (2023).

57. Sasse, A. et al. Benchmarking of deep neural networks for predicting personal gene expression from DNA sequence highlights shortcomings. Nat. Genet. 55, 2060–2064, 10.1038/s41588-023-01524-6 (2023).

58. Tang, Z., Toneyan, S. & Koo, P. K. Current approaches to genomic deep learning struggle to fully capture human genetic variation. Nat. Genet. 55, 2021–2022, 10.1038/s41588-023-01517-5 (2023).

59. Brixi, G. et al. Genome modeling and design across all domains of life with evo 2. .

60. All of Us Research Program Investigators et al. The “All of Us” Research Program. The New Engl. J. Medicine 381, 668–676, 10.1056/NEJMsr1809937 (2019).

61. Bycroft, C. et al. The UK Biobank resource with deep phenotyping and genomic data. Nature 562, 203–209, 10.1038/s41586-018-0579-z (2018). Publisher: Nature Publishing Group.

62. Laub, D. et al. GenVarLoader: An accelerated dataloader for applying deep learning to personalized genomics, 10.1101/2025.01.15.633240 (2025).

63. Laub, D. d-laub/genoray (2025).

64. Morales, J. et al. A joint NCBI and EMBL-EBI transcript set for clinical genomics and research. Nature 604, 310–315, 10.1038/s41586-022-04558-8 (2022).

65. Shannon, C. E. A mathematical theory of communication. The Bell Syst. Tech. J. 27, 379–423, 10.1002/j.1538-7305.1948.tb01338.x (1948).

66. Sundararajan, M., Taly, A. & Yan, Q. Axiomatic attribution for deep networks, 10.48550/arXiv.1703.01365 (2017). 1703.01365[cs].

67. Seitz, E. E., McCandlish, D. M., Kinney, J. B. & Koo, P. K. Interpreting cis-regulatory mechanisms from genomic deep neural networks using surrogate models. Nat. Mach. Intell. 6, 701–713, 10.1038/s42256-024-00851-5 (2024).

68. Tareen, A. & Kinney, J. B. Logomaker: beautiful sequence logos in python. Bioinformatics 36, 2272–2274, 10.1093/bioinformatics/btz921 (2020).

69. Mirdita, M. et al. ColabFold: making protein folding accessible to all. Nat. Methods 19, 679–682, 10.1038/s41592-022-01488-1 (2022). Publisher: Nature Publishing Group.

