## Supplementary Materials for "Genetic background shapes AI-predicted variant effects"

**References** **16**

### 1 Supplementary Tables

| Group | Parameters | References |
|---|---|---|
| ESM1 | 43M, 670M | Meier et al. (2021)[1] |
| ESM1v | 650M | Meier et al. (2021)[1] |
| ESM1b | 650M | Rives et al. (2021)[2] |
| ESM2 | 35M, 650M, 15B | Lin et al. (2023)[3] |
| ESM3 | 1.4B | Hayes et al. (2025)[4] |
| ESMC | 600M | Hayes et al. (2025)[4] |
| AlphaFold2 | 93M | Jumper et al. (2021)[5] |
| Flashzoi | 197M | Hingerl et al. (2024)[6] |
| SpliceAI | 699K | Jaganathan et al. (2019)[7] |

**Supplementary Table 1.** Protein and DNA sequence models used in this study. Rows with $>1$ entry in the *Parameters* column indicates that multiple model sizes were used.

#### 2 Supplementary Figures

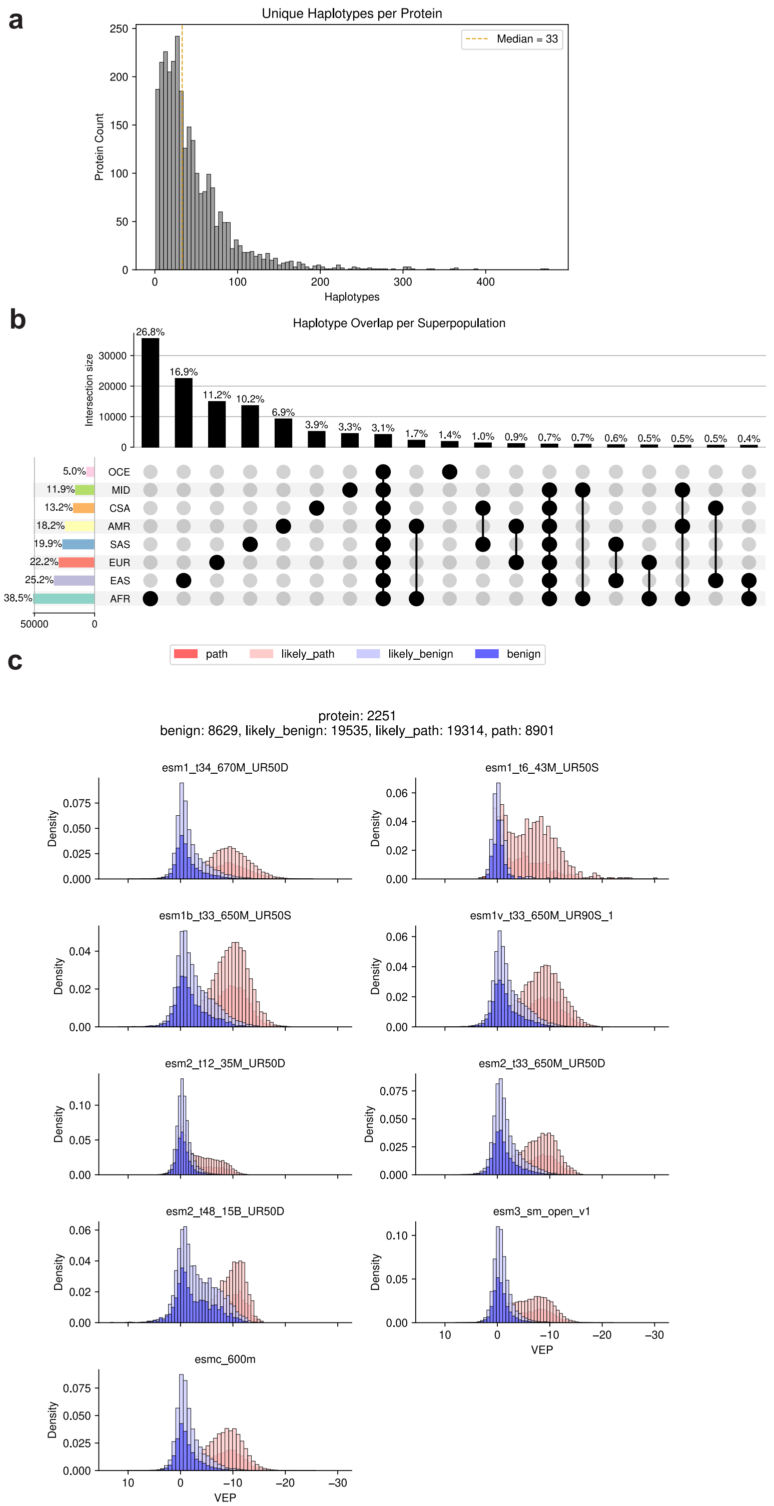

**Supplementary Fig. 1.** **Diverse, population-scale genomes capture protein sequence variation. a**, Histogram of the number of unique haplotypes per protein across all 2,847 proteins in the ProteinGym missense dataset. **b**, An upset plot showing the proportion of haplotypes present in each superpopulation. AFR=African, AMR=Admixed American, CSA=Central South Asian, EAS=East Asian, EUR=European, MID=Middle Eastern, OCE=Oceanian, SAS=South Asian. **c**, VEP distribution histograms stratified by ClinVar pathogenicity labels and faceted by protein language model used to generated the predictions.

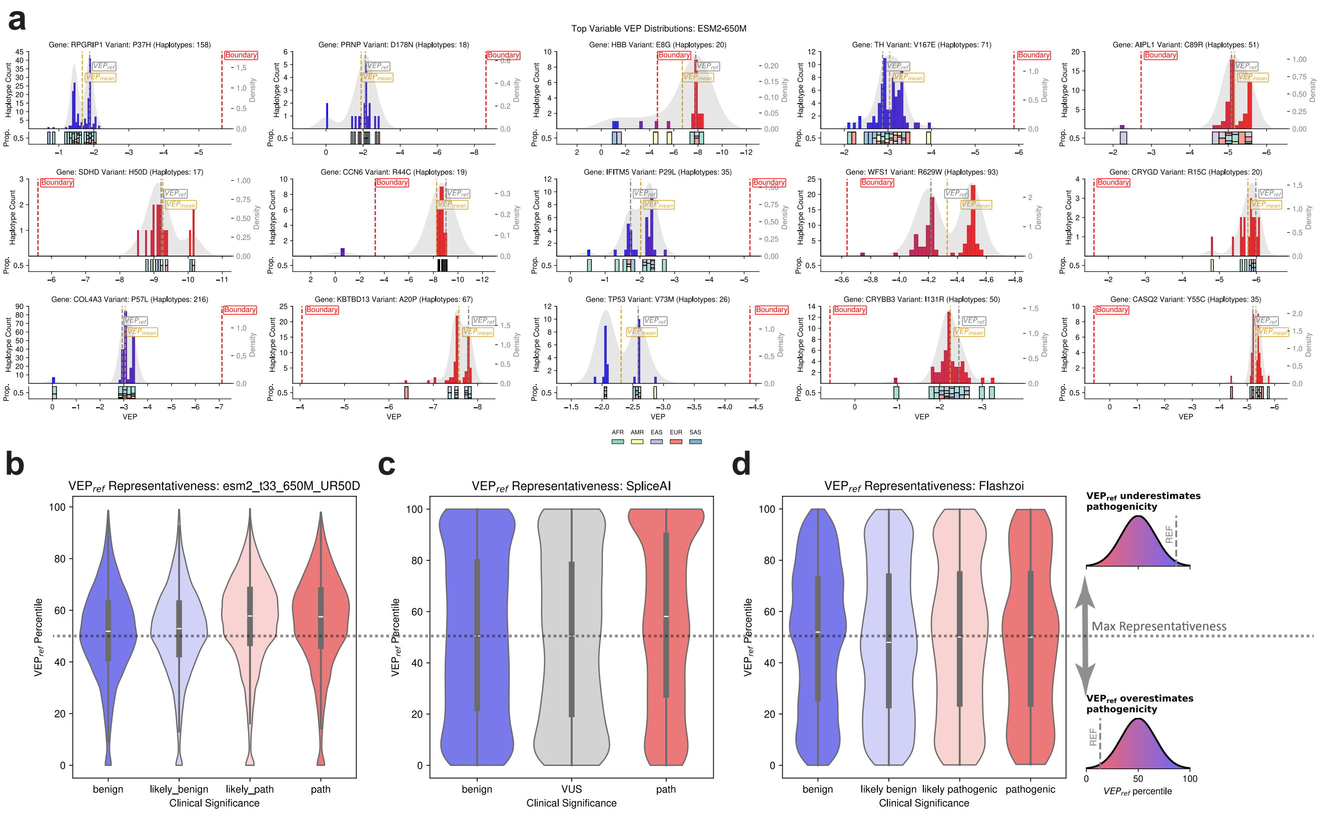

**Supplementary Fig. 2. Reference genome-derived VEP scores are often not representative of full VEP distributions**. **a**, Histograms of ESM2-650M VEP distributions for missense clinical variants with the greatest variability (median absolute deviation). The gold flag indicates the average VEP value across all haplotypes ($VEP_{mean}$), the gray flag indicates the reference-derived VEP ($VEP_{ref}$, and the red flag indicates the benign/pathogenic boundary as empirically inferred from a series of gene-specific Gaussian Mixture models ($Boundary$). **b**, Density distribution of $VEP_{ref}$ percentiles for VEP scores generated with ESM2-650M, stratified by clinical significance annotations from ClinVar. Values above 50% indicate the reference-derived VEP ($VEP_{ref}$) overestimates clinical variant pathogenicity, while values below 50% indicate an underestimation of pathogenicity. A percentile of exactly 50 indicates that $VEP_{ref}$ is maximally representative of the full distribution under this framework. **c**, Distribution of $VEP_{ref}$ percentiles for VEP scores generated with SpliceAI. **d**, Distribution of $VEP_{ref}$ percentiles for VEP scores generated with Flashzoi.

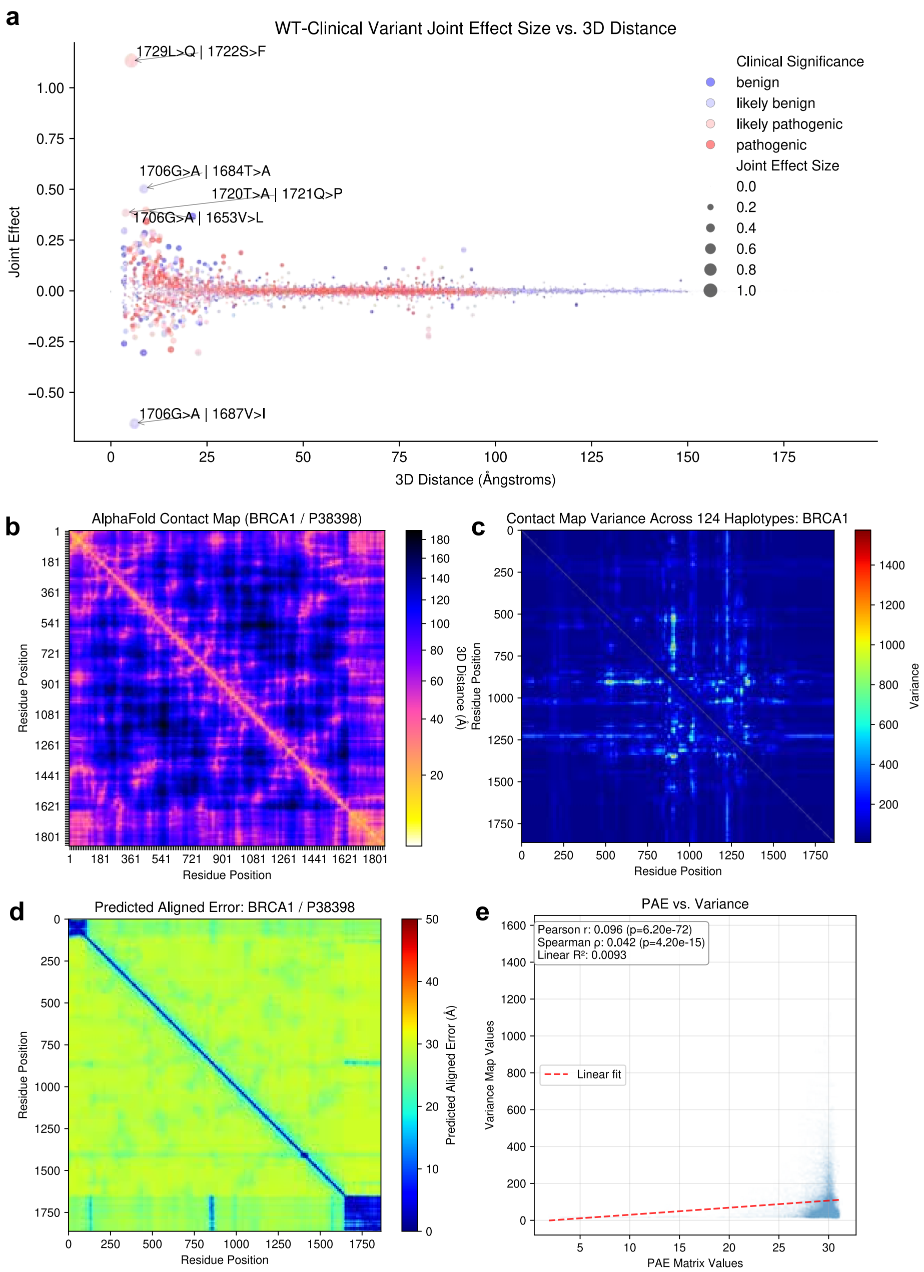

**Supplementary Fig. 3.** **Error in protein folding predictions do not explain inter-haplotype variation in BRCA1. a**, Scatter plot of joint effect sizes of WT and clinical variant pairs (as inferred from surrogate modeling) plotted against 3D residue distance (Å); top five pairs labeled ”WT variant | Clinical variant”). **b**, AlphaFold2 (AF2) distance map for BRCA1 protein using the human reference sequence (Å=Ångstroms). **c**, Variance in AF2 personalized distance maps from 124 unique BRCA1 haplotypes. **d** Predicted Aligned Error (PAE) for AF2 on BRCA reference sequence. **e**, There is a very weak correlation between variance in personalized contacts map and AF2 PAE (Pearson’s $r = 0.098$, $p < 5 \times 10^{-324}$, $n =$ 3,468,906 contact pairs), suggesting that the inter-haplotype variance observed cannot be sufficiently explained by error in AF2’s predictions.

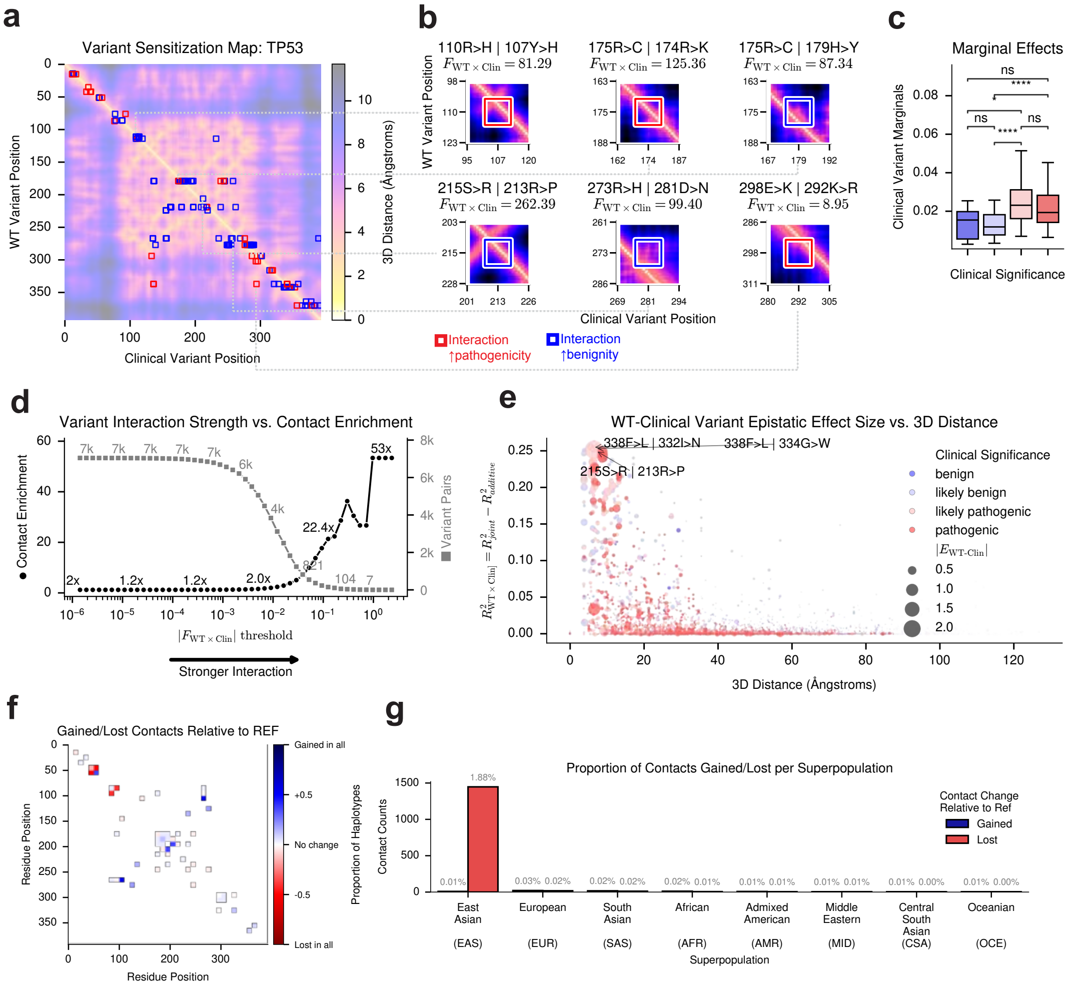

**Supplementary Fig. 4. Background variants interact with clinical variants at 3D protein contact points to influence pathogenicity in TP53 a,** AlphaFold2 (AF2) 3D distance map (in Ångstroms) predicted using the reference protein sequence for TP53., overlaid with variant sensitization map (*red*: background variant ↑ pathogenicity of clinical variant, *blue*: background variant ↑ benignity of clinical variant). **b**, Zoomed-in view of the top 6 WT-Clinical variant pairs with the interactive effects, as inferred from the surrogate model. **c,** Marginal effects from the surrogate model are shown per clinical variant (*y-axis*) and grouped by clinical variant pathogenicity (*x-axis*). Pathogenic variants have significantly higher interaction scores than benign variants (ns: non-significant at $p \geq .05$; $**** : p \leq 0.0001$). **d,** There is a clear positive relationship between the strength of WT-clinical variant joint effects and enrichment for AF2 residue-residue contacts. At each joint effect size threshold, enrichment was computed as observed probability over the expected probability. **e,** Scatter plot showing the relationship between surrogate model joint effect sizes and residue-residue 3D distance (Å). The top three strongest joint effects are labeled with: "WT variant | Clinical variant". **f**, The proportion of population-derived haplotypes that gained (blue) or lost (red) a contact in the reference contact map. All contact maps were first binarized by defining any residue-residue pairs <8 Ångstroms away in the AF2 predictions as a contact (0=no contact, 1=contact). **g,** The number (and percentage) of contacts in the reference contact map that were gained or lost in weighted population-specific contact map. East Asian haplotypes display a significantly larger number of lost contacts relative to the other populations (1.88% vs. <0.03% in all others; 195-fold increase; z-score = 258.75)

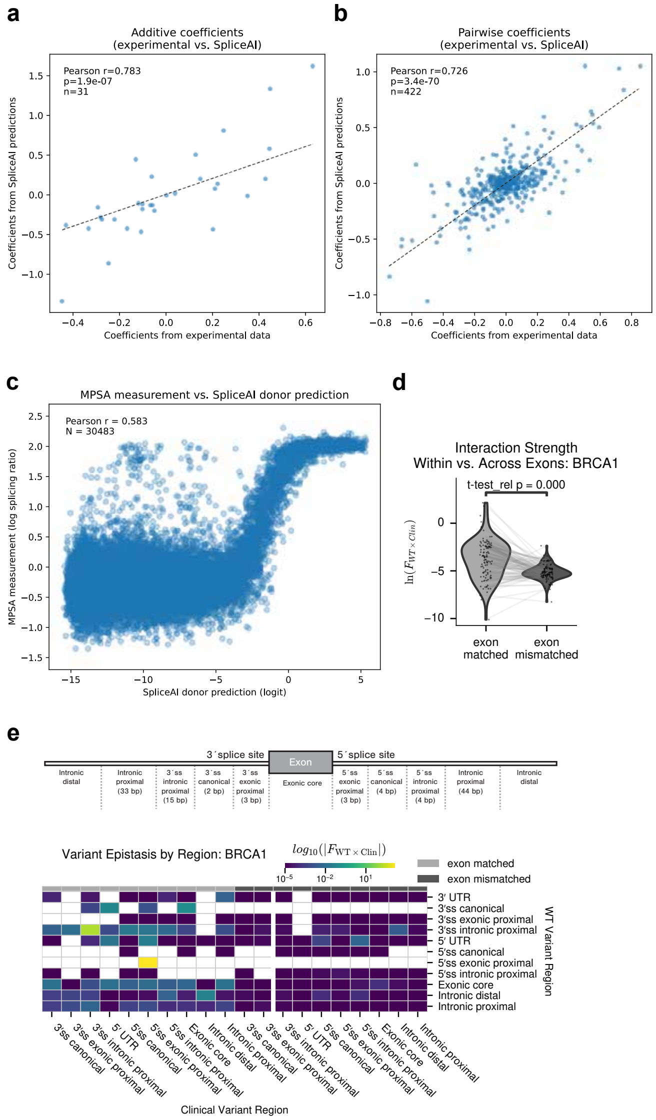

**Supplementary Fig. 5. SpliceAI captures pairwise variant interactions.** Scatter plots comparing coefficients inferred by MAVE-NN linear models fit to experimental massively parallel splicing assay (MPSA) measurements and to SpliceAI-predicted donor logits on the same sequences. **a**, Additive coefficients are correlated between the two models (*x*-axis, MPSA; *y*-axis, SpliceAI; $r = 0.783$, $p = 1.9 \times 10^{-7}$, $n = 31$). **b**, Pairwise coefficients are also correlated (*x*-axis, MPSA; *y*-axis, SpliceAI; $r = 0.726$, $p = 3.4 \times 10^{-70}$, $n = 422$). These results indicate that SpliceAI recapitulates both additive and pairwise epistatic effects underlying splicing regulation. **c**, Correlation between MPSA-measured exon inclusion levels (log splicing ratios) and SpliceAI predicted donor logits for $\sim$30,000 variants at the 9-nt core $5'$ splice site of a *BRCA2* minigene. SpliceAI predictions correlated with experimental measurements (Pearson $r = 0.583$, $N = 30{,}483$). **d**, Mean joint effect size between pairs of *BRCA1* WT and clinical variants as inferred by surrogate modeling of population-derived VEP scores. Variant pairs within the same exon showed stronger interactions than those across different exons (two-sided Mann-Whitney $U$ tests: $p < 0.0001$). **e**, *Top*: Cartoon of annotated splicing regions. *Bottom*: Heatmap of mean interaction strength ($\overline{|Fstat_{\mathrm{WT}\times\mathrm{Clin}}|}$) between WT (*y-axis*) and clinical variants (*x-axis)* in *BRCA1*, aggregated by overlap with splicing annotations (*tick labels*) and exon co-localization (*grayscale bars above*).

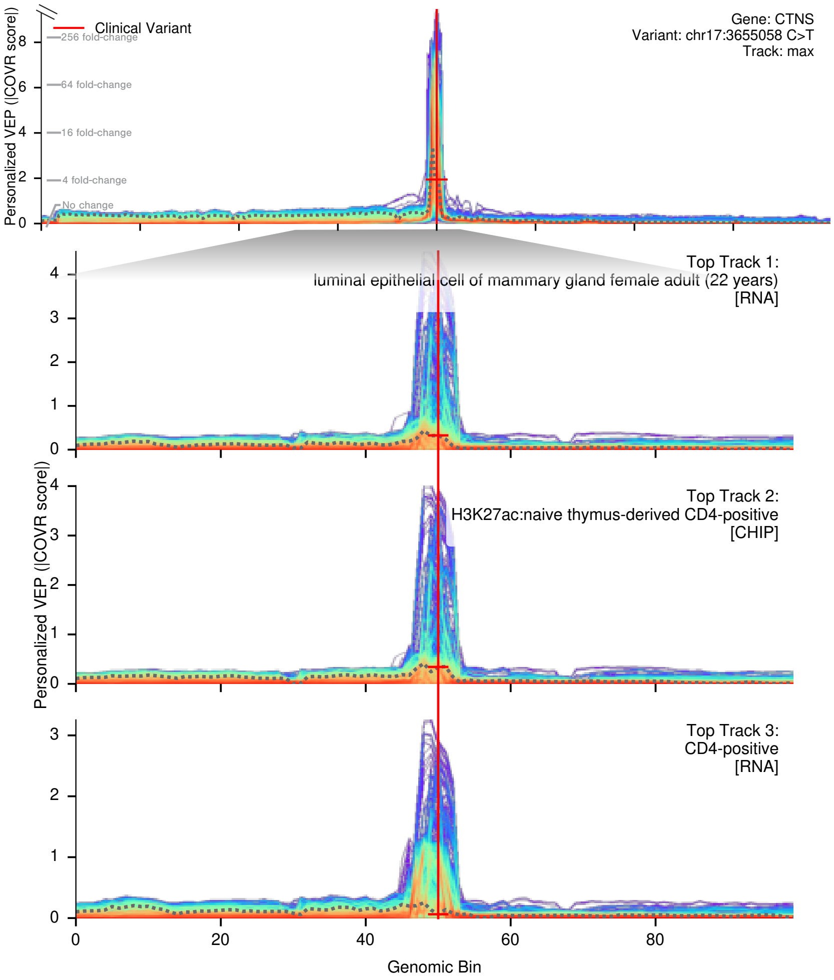

**Supplementary Fig. 6. Flashzoi captures inter-haplotype variation in UTR clinical variants**. To investigate the regulatory basis of this variability, we examined Flashzoi's track–specific outputs for the *CTNS* variant. The tracks with the highest inter-haplotype entropy included RNA expression in mammary and CD4+ tissues, as well as acetylation signals in CD4+ thymus-derived cells. These predictions showed pronounced divergence in response to this variant across individuals; from 0 up to 16-fold changes. This suggests that background variants can substantially modulate gene regulation. Coverage Ratios (COVR) for the pathogenic variant chr17:3655058 C>T (*red vertical line*), with COVR across all tracks shown in top row as the log-scale density (*color*) of haplotypes at each genomic bin (*x-axis*) and VEP value (*y-axis*). The three tracks with the greatest COVR entropy across haplotypes. The reference-derived COVR is shown as a gray dashed line.

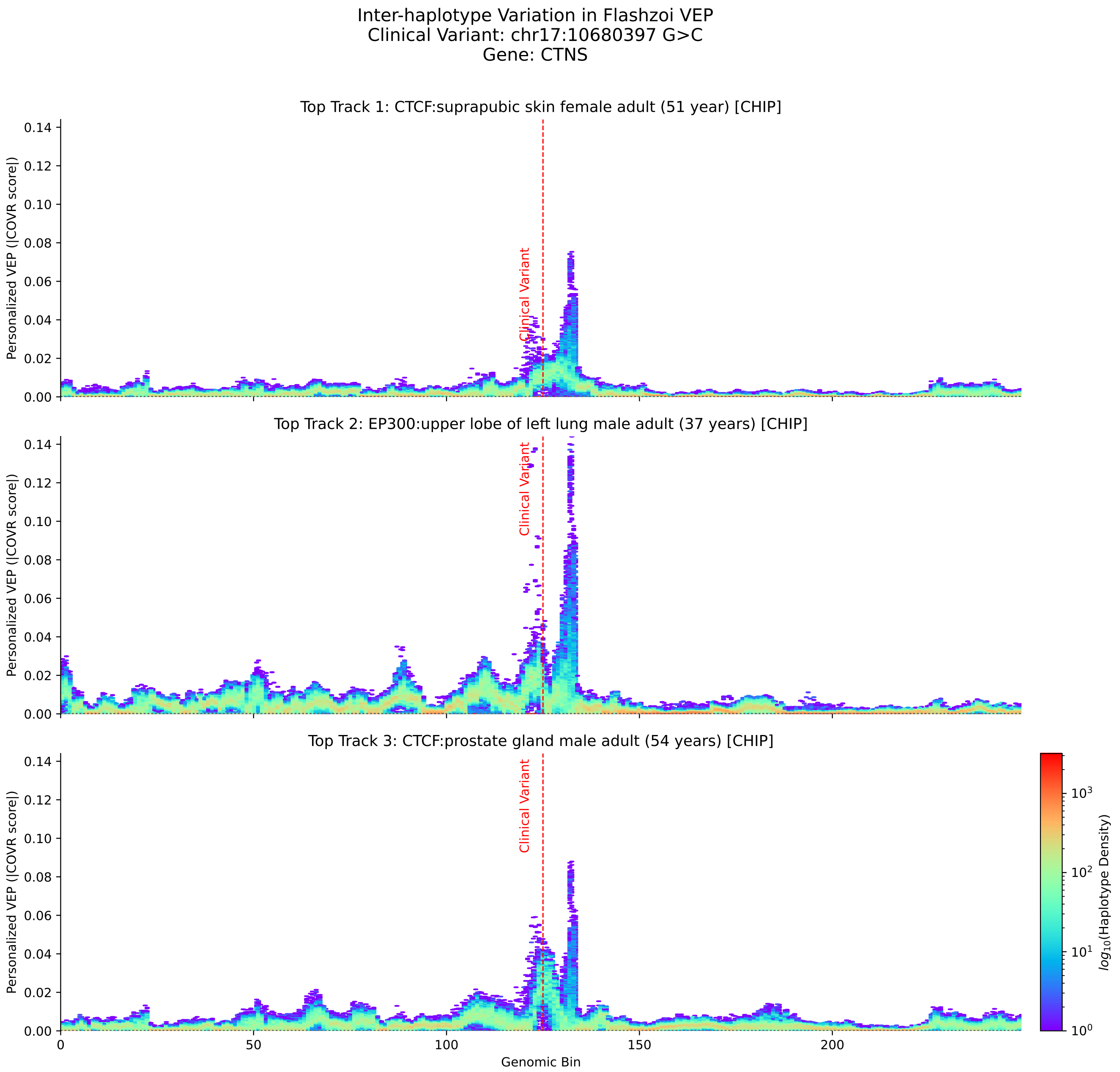

**Supplementary Fig. 7. Flashzoi captures inter-haplotype variation in UTR clinical variants**. For a given 524 kb-wide DNA sequence, we used Flashzoi to predict the read coverage ratio (COVR) of the sequence with an without a given clinical UTR variant, producing thousands of different assay- and tissue-specific track predictions. For each clinical UTR variant, we repeated these predictions across thousands of personalized haplotype sequences. Here, we show the top three tracks (*one per row*) with the greatest inter-haplotype entropy in COVR scores for the variant chr1710680397 G>C, where greater score density corresponds to hotter colors. The red vertical lines indicates the position of the injected clinical variant (chr1710680397 G>C).

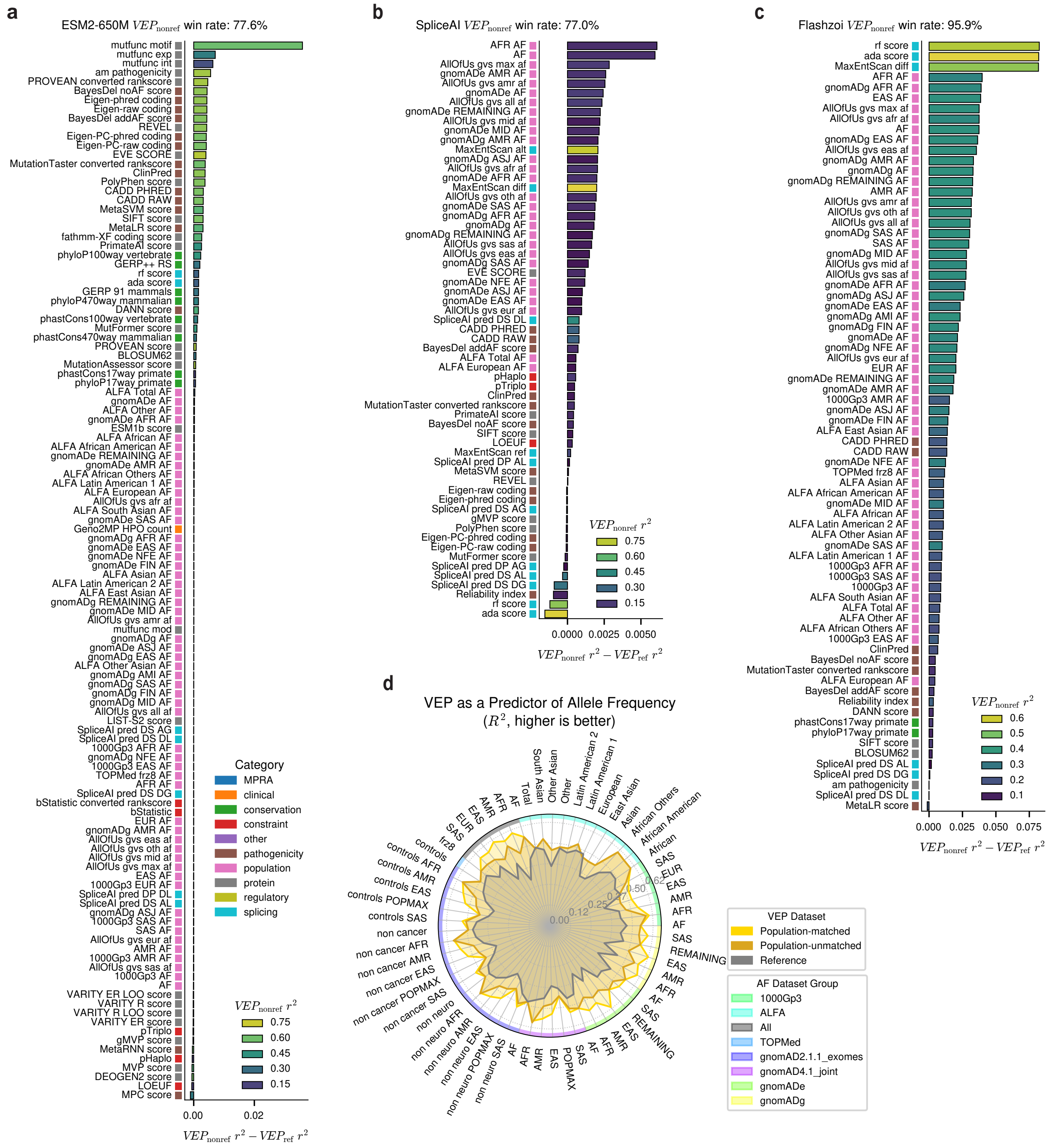

**Supplementary Fig. 8. Population-averaged variant effect predictions improve correlations with relevant Ensembl VEP annotations.** VEP scores derived from only reference sequences ($VEP_{\text{ref}}$) or only population-averaged non-reference sequences ($VEP_{\text{nonref}}$) were correlated with quantitative, genome-wide annotations from Ensembl Variant Effect Predictor (Ensembl VEP)[8]. The difference in Spearman correlation coefficients using $VEP_{\text{nonref}}$ and $VEP_{\text{ref}}$ ($r^2_{\text{nonref}} - r^2_{\text{ref}}$) was then sorted and plotted as bars (*colored by* $r^2_{\text{nonref}}$). Ensembl annotation categories are indicated by the colored boxes to the left of the *y-axis*. This procedure was repeated separately for each sequence model's predictions. Only annotations for which both $VEP_{\text{ref}}$ and $VEP_{\text{nonref}}$ were significantly correlated ($p < 0.05$) are included. **a**, Correlations between ESM2 VEP scores and variant annotations. **b**, Correlations between Flashzoi VEP scores and variant annotations. **c**, Correlations between SpliceAI VEP scores and variant annotations. **d**, Superpopulation-averaged VEP scores were used to predict allele frequency from matched populations, which consistently outperformed both population-agnostic VEP ($VEP_{\text{mean}}$) and reference-only VEP ($VEP_{\text{ref}}$).

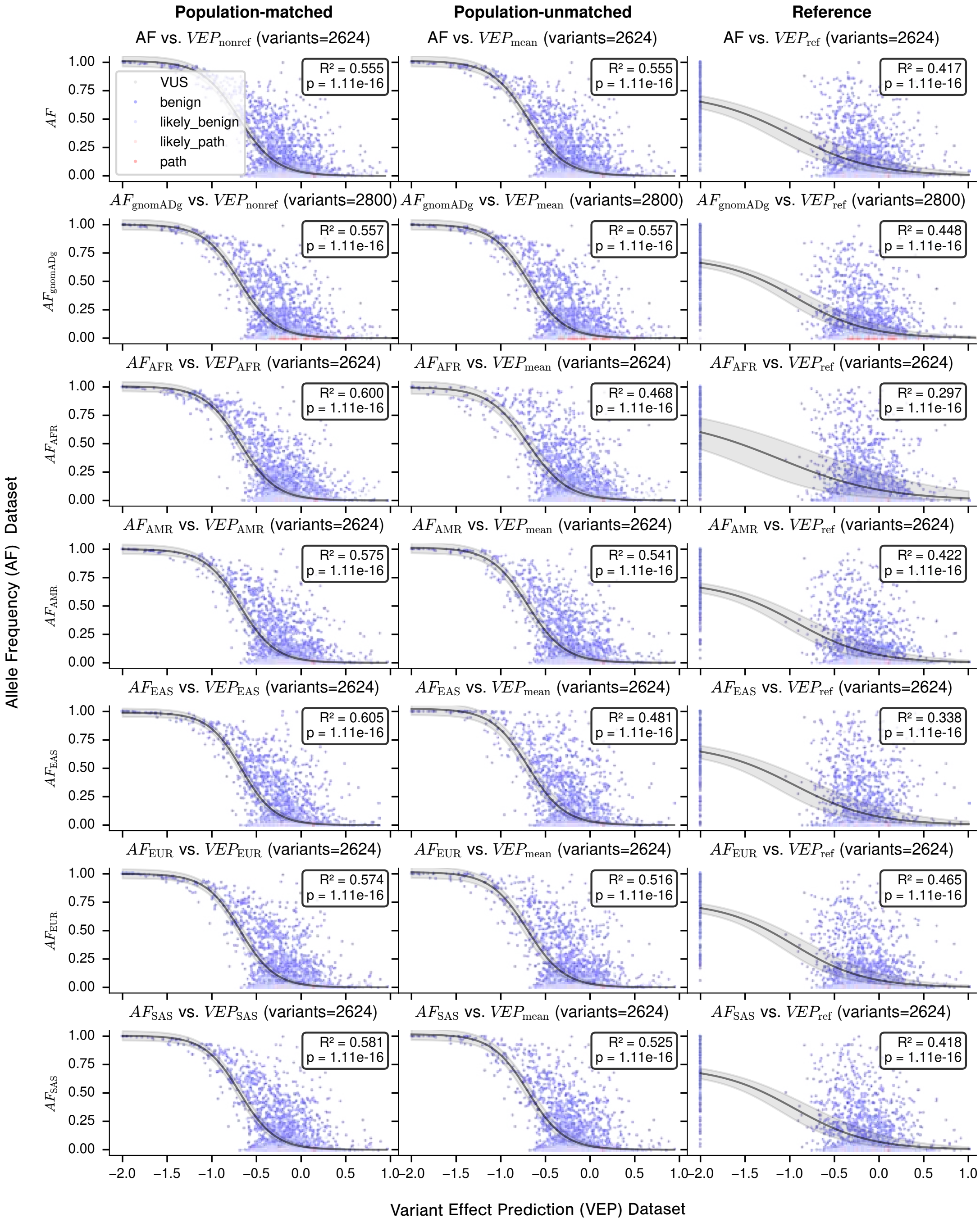

**Supplementary Fig. 9. Personalized VEP scores outperform reference VEP scores in predicting allele frequencies, especially when matched by superpopulation**. Per-variant VEP Flashzoi scores were averaged across all samples ($VEP_{mean}$) , all samples except for the reference ($VEP_{\text{nonref}}$), only the reference ($VEP_{\text{ref}}$), and superpopulation-specific samples only ($VEP_{\text{AFR}}$=African, $VEP_{\text{AMR}}$=Admixed American, $VEP_{\text{EAS}}$=East Asian, $VEP_{EUR}$=European, $VEP_{SAS}$=South Asian). Then, each allele frequency (AF) annotation was correlated with a superpopulation-matched VEP ($VEP_{[match]}$, *first column*), VEP averaged across all populations without matching ($VEP_{\text{mean}}$, *second column*), or reference-derived VEP ($VEP_{\text{ref}}$, *third column*). Model fit consistently improved from: $VEP_{[matched]} > VEP_{mean} >> VEP_{ref}$

### 3 Supplementary Methods

#### 3.1 Linear Model for Joint Effect Estimation

To quantify the joint effects of wild-type (WT) variants and clinical variants on variant effect predictor (VEP) scores, we used regularized linear regression via `sklearn.linear_model.Ridge` (L2 regularization). For each gene, we constructed a binary matrix $\mathbf{X}$ (#haplotypes $\times$ #WT variants), representing the presence or absence of each WT variant in each haplotype, and a continuous matrix $\mathbf{Y}$ (#haplotypes $\times$ #sites) containing VEP scores for each haplotype-site pair. We then fit a multi-target ridge regression model:

$$\mathbf{Y} = \mathbf{X}\beta + \varepsilon$$

Here, $\beta$ (a WT variants $\times$ sites coefficient matrix, as returned by `Ridge.coef_`) is fit by minimizing squared error plus a regularization penalty established by the strength parameter $\alpha$. The absolute value of each coefficient, $|\beta_{ij}|$, measures the magnitude of the joint effect between WT variant $i$ and site $j$, and the sign of $\beta_{ij}$ encodes the direction of the effect (negative values indicating more pathogenic, positive indicating more benign).

For each WT variant-site pair, we computed:

1. Absolute joint effect magnitude $|\beta_{ij}|$
2. Signed joint effect $\beta_{ij}$
3. The number of haplotypes containing the WT variant (computed by summing the corresponding binary indicator column)
4. Weighted joint effect: $\sqrt{|\beta_{ij}| \times n_{\text{haplotypes}}}$ (to balance effect size and sample count)

Model quality was evaluated using standard metrics from `sklearn.metrics`: $R^2$, explained variance, mean squared error (MSE), root mean squared error (RMSE), and mean absolute error (MAE).

#### 3.2 Interaction Testing

To identify non-additive interactions (as opposed to additive effects), we implemented a statistical testing procedure that directly compares observed joint effects to the sum of their separate effects.

For each WT variant $i$ and clinical variant $j$ pair, we first computed:

1. **Individual WT variant effect:** The mean VEP score difference between haplotypes where WT variant $i$ is present (WT=1) vs. absent (WT=0), averaging across all clinical variants:

   $$\mathrm{E}_{\mathrm{WT},i} = \frac{1}{|\mathscr{S}|} \sum_{s \in \mathscr{S}} \left( \overline{\mathrm{VEP}}(\mathrm{WT}_i{=}1, s) - \overline{\mathrm{VEP}}(\mathrm{WT}_i{=}0, s) \right)$$

   where $\mathscr{S}$ is the set of all clinical variant sites.

2. **Individual clinical variant effect:** The average VEP score for clinical variant $j$ relative to the overall VEP baseline (mean across all haplotypes and all clinical variants):

   $$\mathrm{E}_{\mathrm{Clin},j} = \overline{\mathrm{VEP}}(\mathrm{Clinical}_j) - \overline{\mathrm{VEP}}_{\mathrm{baseline}}$$

The **expected additive effect** is then:

$$\mathrm{E}_{\mathrm{additive}} = \mathrm{E}_{\mathrm{WT},i} + \mathrm{E}_{\mathrm{Clin},j}$$

The **observed joint effect** is given by the corresponding $\beta_{ij}$ coefficient from the main `Ridge` model, which reflects the effect when both WT variant $i$ and clinical variant $j$ are present. To statistically assess whether this joint effect is interactive, we compared two nested models, both fit via `sklearn.linear_model.Ridge`:

- **Additive model:**

  $$\mathrm{VEP} = \beta_0 + \beta_1 \cdot \mathrm{WT}$$

  where the WT variant's effect is constant (clinical variant is absorbed in the intercept).

- **Interaction model:**

$$\text{VEP} = \beta_0 + \beta_1 \cdot \text{WT} + \beta_2 \cdot (\text{WT} \times \Delta_{\text{deviation}})$$

where $\Delta_{\text{deviation}} = \text{E}_{\text{additive}} - \text{E}_{\text{WT},i}$, capturing deviation from pure additivity, with $\beta_2$ representing the interaction strength.

Model fits for both cases were obtained using `Ridge` (with the same regularization setup), and the null hypothesis ($\beta_2 = 0$, i.e., no interaction) was tested via an F-test comparing the residual sum of squares (RSS):

$$F = \frac{(RSS_{\text{additive}} - RSS_{\text{interaction}})/(df_{\text{additive}} - df_{\text{interaction}})}{RSS_{\text{interaction}}/df_{\text{interaction}}}$$

where $df_{\text{additive}} = n-2$ and $df_{\text{interaction}} = n-3$ ($n =$ number of haplotypes). P-values were calculated using `scipy.stats.f.cdf` (F-distribution, 1 and $df_{\text{interaction}}$ degrees of freedom). Since `Ridge` is a regularized model, degrees of freedom are approximate, but the F-test remains valid for comparing nested models.

A pair was classified as interactive if the interaction model fit led to a statistically significant improvement ($p < 0.05$ by default, controlled via the `interaction_pvalue_threshold` parameter) over the additive-only model. For each test, we recorded:

1. Interactive $p$-value and F-statistic
2. $R^2$ and MSE for both the additive and interaction models
3. $R^2$ improvement ($\Delta R^2 = R^2_{\text{interaction}} - R^2_{\text{additive}}$)
4. Interaction coefficient $\beta_2$ (strength of interaction)
5. The joint effect $\beta_{ij}$, expected additive effect, and their deviation
6. Individual WT variant and clinical variant effects

To ensure robust inference, we restricted interaction tests to variant pairs meeting the following criteria:

1. At least 10 haplotypes with complete, non-missing data for both the WT variant and the clinical variant site
2. Sufficient variation in WT variant status (std $> 10^{-10}$)
3. Both WT=0 and WT=1 represented in the dataset

These filters prevent false positives from insufficient or uninformative data.

##### 3.3 Window-Based Interaction Testing for Site-Centered Models

For analyses where clinical variants are analyzed individually with site-centered genomic windows (e.g., SpliceAI predictions), we implemented an alternative interaction testing approach that accounts for the fact that each clinical variant is modeled separately with a distinct set of WT variants within its genomic window.

**Model Training Strategy** Rather than fitting a single multi-target model across all clinical variants simultaneously, we fit separate Ridge regression models for each clinical variant $j$:

$$\mathbf{Y}_j = \mathbf{X}_j \beta_j + \varepsilon_j$$

where $\mathbf{X}_j$ is a binary matrix (#haplotypes $\times$ #WT variants within window $j$) containing only WT variants within a genomic window centered on clinical variant $j$ (typically $\pm 5$ kb), and $\mathbf{Y}_j$ is a vector (#haplotypes $\times$ 1) containing VEP scores for clinical variant $j$ only. This window-based approach ensures that each model focuses on WT variants that were actually seen by the sequence model during VEP inference time.

**Cross-Model WT Effect Estimation** A key difference from the multi-target approach is how individual WT variant effects are computed. Since each model includes a different set of WT variants (determined by the genomic window), we cannot reliably compute WT individual effects within a single model context. Instead, we compute the individual effect of each WT variant $i$ by averaging its effect across *all* clinical variant surrogate models where it appears:

$$\text{Effect}_{\text{WT},i} = \frac{1}{|\mathscr{M}_i|} \sum_{m \in \mathscr{M}_i} \left( \overline{\text{VEP}}(\text{WT}_i{=}1, m) - \overline{\text{VEP}}(\text{WT}_i{=}0, m) \right)$$

where $\mathscr{M}_i$ is the set of all surrogate models(indexed by $m$) that include WT variant $i$ in their genomic window. This cross-model averaging provides a robust estimate of the WT variant's average effect across different genomic contexts, which serves as the expected additive effect when testing for interactions.

**Interaction Testing Procedure** For each WT variant $i$ and clinical variant $j$ pair, interaction testing proceeds as follows:

1. **Expected additive effect:** Since we are testing one clinical variant at a time, the clinical variant effect is constant across all haplotypes (absorbed into the intercept). Therefore, the expected additive effect is simply:

   $$\text{Effect}_{\text{additive}} = \text{Effect}_{\text{WT},i}$$

   where $\text{Effect}_{\text{WT},i}$ is computed across all models as described above.

2. **Observed joint effect:** The coefficient $\beta_{ij}$ from the model for clinical variant $j$ represents the observed joint effect.

3. **Model comparison:** We fit the same additive and interaction models as in the multi-target approach:

   - **Additive model:** $\text{VEP}_j = \beta_0 + \beta_1 \cdot \text{WT}_i$
   - **Interaction model:** $\text{VEP}_j = \beta_0 + \beta_1 \cdot \text{WT}_i + \beta_2 \cdot (\text{WT}_i \times \Delta_{\text{deviation}})$

   where $\Delta_{\text{deviation}} = \beta_{ij} - \text{Effect}_{\text{WT},i}$.

4. **Statistical testing:** The F-test and p-value calculation proceed identically to the multi-target approach.

**Implementation Details and Computational Workflow** All analyses were conducted in Python 3.12.9, and all core methods have been implemented as custom functions available in the public GitHub repository:
https://github.com/bschilder/VEP_DNA

Linear regression models for both additive effects and interaction testing use `sklearn.linear_model.Ridge`, statistical hypothesis tests rely on `scipy.stats.f`, and data manipulation leverages `pandas`.

The workflow is structured in a flexible two-stage procedure that works for both the multi-target and window-based approaches:

1. **Model Training:** First, the main linear models are fit for either all clinical variants simultaneously (multi-target mode) or for each clinical variant individually in their respective genomic windows (window-based mode). This is done using the `wtvariants_to_vep_linear_model` function, which supports Ridge regression, user-specified regularization, and (if desired) integrated interaction testing via the `test_epistasis` flag.

2. **Interaction Testing:** After all models are trained, interactivity is assessed using the `test_epistasis_across_models` function. This computes WT individual effects by aggregating across all relevant clinical variant models—enabling the calculation of expected additive effects even for window-based approaches where each model contains a different set of WT variants. Batch interaction testing is efficiently performed across all possible model-variant pairs, handling scenarios where variant sets differ due to genomic windowing.

The above design enables:

- Aggregation and estimation of WT individual effects across all models, ensuring robust expected effect calculations for use in interaction testing.
- Scalable, efficient batch processing of interaction tests, regardless of whether a multi-target or window-based modeling approach is used.
- Generation of a combined dataframe that contains both the initial interaction results and all interaction annotations and statistics. Updated model dictionaries are returned with integrated interaction results.

##### 3.4 Calculating contact enrichment for WT-clinical variant joint effects

Let $L$ denote the protein length (number of residues). We define the binary contact map

$$C \in \{0,1\}^{L\times L},$$

where

$$C_{ij} = \begin{cases} 1 & \text{if residues } i \text{ and } j \text{ are in physical contact,} \\ 0 & \text{otherwise.} \end{cases}$$

with $C_{ii} = 0$ for all $i$.

The total number of possible residue-residue pairs is

$$N_{\text{total}} = L(L-1).$$

Let

$$Z \in \mathbb{R}^{L\times L}$$

denote the matrix of absolute joint effect scores for each residue pair.

For a given threshold $t \geq 0$, we define the binary indicator of significant joint effects as

$$X_{ij}^{(t)} = \begin{cases} 1 & \text{if } Z_{ij} > t, \\ 0 & \text{otherwise.} \end{cases}$$

The number of significant joint effects at threshold $t$ is

$$N_{\text{sig}}(t) = \sum_{i\neq j} X_{ij}^{(t)},$$

the number of predicted contacts is

$$N_{\text{contacts}} = \sum_{i\neq j} C_{ij},$$

and the observed overlap is

$$N_{\text{overlap}}(t) = \sum_{i\neq j} X_{ij}^{(t)} \cdot C_{ij}.$$

The observed probability of overlap at threshold $t$ is

$$\hat{p}_{\text{obs}}(t) = \frac{N_{\text{overlap}}(t)}{N_{\text{sig}}(t)}.$$

The expected probability under the null model is independent of $t$:

$$p_{\text{exp}} = \frac{N_{\text{contacts}}}{N_{\text{total}}} = \frac{N_{\text{contacts}}}{L(L-1)}.$$

For each threshold $t$, significance is assessed via a binomial test:

$$p\text{-value}(t) = \Pr\left(\text{Binomial}\big(n = N_{\text{sig}}(t),\, p = p_{\text{exp}}\big) \geq N_{\text{overlap}}(t)\right) \quad \text{(two-sided).}$$

Finally, the fold enrichment at threshold $t$ is defined as

$$E(t) = \frac{\hat{p}_{\text{obs}}(t)}{p_{\text{exp}}}.$$

$$\begin{cases} E(t) > 1 & \text{enrichment: contacts occur more often than expected,} \\ E(t) < 1 & \text{depletion: contacts occur less often than expected.} \end{cases}$$

##### 3.5 Testing for relationships between variant effect and frequency

The relationship between VEP scores and allele frequencies (AF) was modeled using a three-parameter logistic function: $f(x) = \frac{L}{1+\exp(-k(x-x_0))}$ , where $L$ is the maximum value, $k$ is the growth rate, and $x_0$ is the midpoint. Parameters were estimated via nonlinear least squares using `scipy.optimize.curve_fit` (maximum function evaluations = 10,000). Model fit was assessed using the coefficient of determination ($R^2$) and an F-test for significance. 95% confidence intervals were computed from the parameter covariance matrix using $\pm 1.96$ standard errors and displayed as shaded regions around the fitted curve.

### 4 Extended Discussion: The Structural Basis for Personalized Variant Effect Prediction

#### 4.1 The Reference Genome as a Fragmented Mosaic

The human reference genome (hg38/GRCh38) is frequently treated as a continuous biological gold standard; however, its architectural origins introduce significant structural biases into variant effect predictions. As detailed by the Genome Reference Consortium[9], hg38 is a "collapsed haploid" mosaic. It was constructed using Bacterial Artificial Chromosome (BAC) clones, which are physical fragments of DNA typically averaging 150kb in length.

This construction method effectively performed a "mechanical phasing" at the bench: each bacterial colony contained only one physical fragment from a donor's genome, either the maternal or the paternal copy. Consequently, the primary assembly represents only one side of a donor's diploid identity at any given locus, with the alternative parental haplotype discarded during assembly. Furthermore, because these 150kb clones were "tiled" together from multiple individuals to form a single linear sequence, the reference genome contains artificial "seams" or junctions where the sequence flips between different donors and different parental strands. This creates a "Frankenstein" haplotype that has never existed in a natural human cell.

#### 4.2 Context-Dependent Model Sensitivity

Our findings demonstrate that personalized variant effect predictor (pVEP) scores are partly dependent on the context window of the predictive model employed. We observed that haplotype context significantly altered effect estimates for large-context models like Flashzoi (250kb), while local-context models like SpliceAI (10kb) remained relatively stable.

This disparity can be explained by the 150kb scale of the reference building blocks:

- **Local Coherence (10kb):** At a 10kb scale, a model is almost certainly looking at a sequence derived from a single BAC clone. At this resolution, the reference is internally "phased" and consistent, even if it only represents one parental copy. Small-window models like SpliceAI are thus less sensitive to the mosaic nature of the reference.
- **Structural Fragmentation (250kb):** In contrast, a 250kb window is statistically likely to span at least one BAC clone boundary. For models like Borzoi/Flashzoi[6,10], the reference sequence is structurally fragmented, potentially placing an enhancer from one donor or parental strand in proximity to a promoter from another. By resolving these distal seams, the pVEP framework restores the long-range cis-regulatory continuity that large-context architectures are designed to interpret.

#### 4.3 Addressing the Limitations of Short-Read WGS

A potential critique of pVEP, as used in our study, is its reliance on short-read whole-genome sequencing (srWGS), which is subject to mapping biases and statistical switch errors. However, our analysis suggests that a statistically phased diploid genome, even with imperfect switch accuracy, remains a more biologically faithful template than the reference mosaic.

While srWGS may contain "noise," the hg38 reference contains "omissions." The reference is missing approximately 300 million base pairs of DNA found in non-European and non-West African populations[9]. By utilizing pVEP in diverse cohorts of individuals, we incorporate these population-specific haplotypes and preserve the diploid nature of human biology. This approach moves the VEP field away from a *one-variant-one-effect* paradigm and toward a framework that recognizes the essential role of genetic background in modulating variant pathogenicity.

#### 4.4 Conclusion and Future Directions

The systematic ignoring of genetic context in conventional reference-centric VEP frameworks is not merely a technical oversight but a substantive equity concern. Our results highlight that the populations least represented in hg38—such as East Asian, South Asian, and Indigenous American groups—stand to gain the most from personalized frameworks. As we transition toward a pangenomic era, the transition from mosaic reference sequences to truly diploid, population-scale genetic backgrounds will be essential for the stable and equitable evaluation of AI-based sequence models.

### References

1. Meier, J. *et al.* Language models enable zero-shot prediction of the effects of mutations on protein function, https://doi.org/10.1101/2021.07.09.450648 (2021).
2. Rives, A. *et al.* Biological structure and function emerge from scaling unsupervised learning to 250 million protein sequences. *Proc. Natl. Acad. Sci.* **118**, e2016239118, https://doi.org/10.1073/pnas.2016239118 (2021).
3. Lin, Z. *et al.* Evolutionary-scale prediction of atomic-level protein structure with a language model. *Science* **379**, 1123–1130, https://doi.org/10.1126/science.ade2574 (2023).
4. Hayes, T. *et al.* Simulating 500 million years of evolution with a language model. *Science* **0**, eads0018, https://doi.org/10.1126/science.ads0018 (2025).

5. Jumper, J. *et al.* Highly accurate protein structure prediction with AlphaFold. *Nature* **596**, 583–589, https://doi.org/10.1038/s41586-021-03819-2 (2021).
6. Hingerl, J. C., Karollus, A. & Gagneur, J. Flashzoi: an enhanced borzoi for accelerated genomic analysis. *Bioinformatics* **41**, btaf467, https://doi.org/10.1093/bioinformatics/btaf467 (2025).
7. Jaganathan, K. *et al.* Predicting splicing from primary sequence with deep learning. *Cell* **176**, 535–548.e24, https://doi.org/10.1016/j.cell.2018.12.015 (2019).
8. McLaren, W. *et al.* The ensembl variant effect predictor. *Genome Biol.* **17**, 122, https://doi.org/10.1186/s13059-016-0974-4 (2016).
9. Schneider, V. A. *et al.* Evaluation of GRCh38 and de novo haploid genome assemblies demonstrates the enduring quality of the reference assembly. *Genome Res.* **27**, 849–864, https://doi.org/10.1101/gr.213611.116 (2017).
10. Linder, J., Srivastava, D., Yuan, H., Agarwal, V. & Kelley, D. R. Predicting RNA-seq coverage from DNA sequence as a unifying model of gene regulation. *Nat. Genet.* 1–13, https://doi.org/10.1038/s41588-024-02053-6 (2025).
